# *Toxoplasma gondii* requires its plant-like heme biosynthesis pathway for infection

**DOI:** 10.1101/753863

**Authors:** Amy Bergmann, Katherine Floyd, Melanie Key, Carly Dameron, Kerrick C. Rees, Daniel C. Whitehead, Iqbal Hamza, Zhicheng Dou

**Affiliations:** Department of Biological Sciences, Clemson University, Clemson, South Carolina, USA 29634; Department of Chemistry, Clemson University, Clemson, South Carolina, USA 29634; Department of Animal and Avian Sciences, University of Maryland, College Park, Maryland, USA 20742; Department of Cell Biology and Molecular Genetics, University of Maryland, College Park, Maryland, USA 20742

## Abstract

Heme, an iron-enclosed organic ring, is essential for virtually all living organisms by serving as a prosthetic group in proteins that function in diverse cellular activities ranging from diatomic gas transport and detection to mitochondrial respiration to detoxification. Cellular heme levels in microbial pathogens can be a composite of endogenous *de novo* synthesis or exogenous uptake of heme or heme synthesis intermediates^1,2^. Intracellular pathogenic microbes switch routes for heme supply when heme availability in their replicative environment fluctuates through infections^2^. Here, we show that the *Toxoplasma gondii*, an obligate intracellular human pathogen, encodes a functional heme biosynthesis pathway. A chloroplast-derived organelle, termed apicoplast, is involved in the heme production. Genetic and chemical manipulation revealed that *de novo* heme production is essential for *T. gondii* intracellular growth and pathogenesis. Surprisingly, the herbicide oxadiazon significantly impaired *Toxoplasma* growth, consistent with phylogenetic analyses that show *T. gondii* protoporphyrinogen oxidase is more closely related to plants than mammals. We further improve upon this inhibition by 15-to 25-fold with two oxadiazon derivatives, providing therapeutic proof that *Toxoplasma* heme biosynthesis is a druggable target. As *T. gondii* has been used to model other apicomplexan parasites^3^, our study underscores the utility of targeting heme biosynthesis in other pathogenic apicomplexans.

---

Human protozoan pathogens share some common nutrient metabolism pathways with their counterparts in the host but show distinct features. Apicomplexan parasites, including *Toxoplasma gondii* and *Plasmodium spp.,* encode a complete *de novo* heme biosynthesis pathway in their genomes^2^ (Fig. 1a). In *Toxoplasma*, a previous study of ectopic overexpression of all 8 heme biosynthetic proteins revealed that these enzymes are delivered to three subcellular locations^4^, including the mitochondrion, cytoplasm, and apicoplast. The apicoplast is a remnant chloroplast and specifically exists in apicomplexan parasites. Moreover, the second enzyme residing within the parasite’s heme biosynthesis pathway, *Toxoplasma* porphobilinogen synthase (TgPBGS), was expressed in an active recombinant form^5,6^. These previous findings revealed that the heme biosynthetic enzymes in *Toxoplasma* carry targeting signals for their subcellular localization, and one of these enzymes, TgPBGS, is enzymatically functional. However, it still remains unknown whether the entire pathway is active for heme production during *Toxoplasma* infections, and this pathway plays a critical role in *Toxoplasma* infection.

**Fig. 1.**
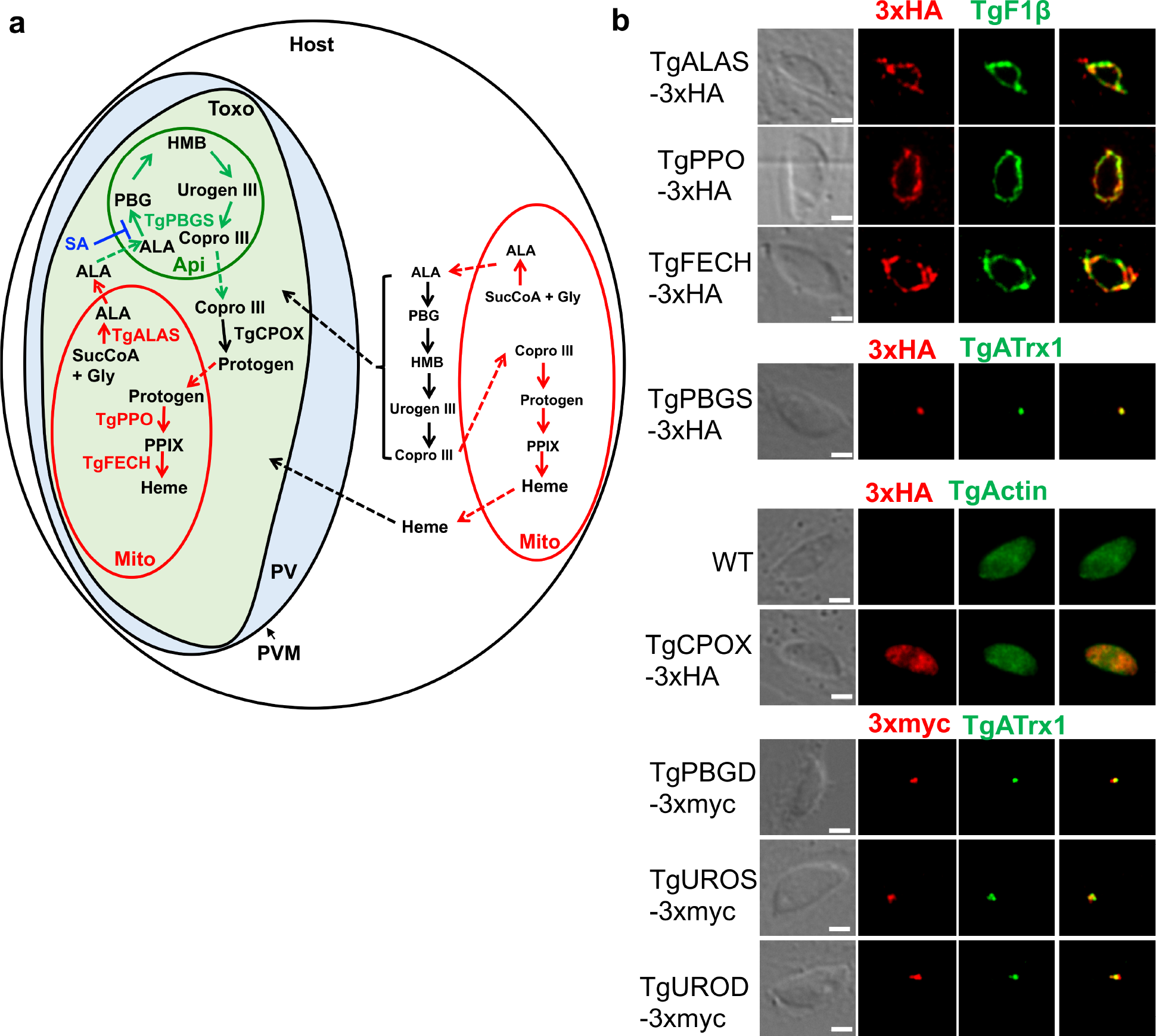
*Toxoplasma gondii* encodes its *de novo* heme biosynthetic pathway within 3 subcellular locations. **a,** The working model of the *de novo* heme biosynthesis in *Toxoplasma* parasites. The enzymes catalyzing the *de novo* heme biosynthesis are distributed within three subcellular locations in the parasites, whereas they are only localized in the mitochondria and cytoplasm in mammals. **b,** Determination of the expression of the heme biosynthetic genes in *Toxoplasma* during its acute infection and their subcellular locations by endogenous gene tagging with 3xHA or 3xmyc epitopes. A subunit of *Toxoplasma* mitochondrial ATPase (TgF1β) and an apicoplast-associated thioredoxin family protein (TgATrx1) were used as the mitochondria and apicoplast markers, respectively. TgActin was used as a cytoplasm marker. Bar = 2 μm. ALA, Aminolevulinic acid; ALAS, Aminolevulinic acid synthase; Api, apicoplast; Copro III, coproporphyrinogen III; CPOX, Coproporphyrinogen III oxidase; FECH, Ferrochelatase; Gly, glycine; HMB, hydroxymethylbilan; IVN, intravacuolar network; Mito, mitochondria; PBG, porphobilinogen; PBGD, Porphobilinogen deaminase; PBGS, Porphobilinogen synthase; PPIX, protoporphyrin IX; PPO, Protoporphyrinogen oxidase; Protogen, protoporphyrinogen IX; PV, parasitophorous vacuole; PVM, parasitophorous vacuole membrane; SucCoA, Succinyl-CoA; Toxo, *Toxoplasma gondii*; UROD, Uroporphyrinogen III decarboxylase; Urogen III, uroporphyrinogen III; UROS, Uroporphyrinogen III Synthase.

As an intracellular parasite, *Toxoplasma* utilizes the host plasma membrane to create its own membrane-bound compartment for intracellular replication, termed the parasitophorous vacuole (PV)^7^. The PV membrane (PVM) is permeable for small solutes. Studies have demonstrated that putative nutrient pores exist on the PVM that allows small substances with molecular weights less than ~1,300 Da to diffuse into the PV^8^. A recent study revealed that the parasites use two dense granule proteins, TgGRA17 and TgGRA23, to form a pore structure on the PVM, which allows small molecules to enter the PV from host cytoplasm^9^. One such small molecule is heme, a prosthetic group found in a number of essential proteins, leading to the hypothesis that parasites can hijack the host’s heme or heme synthesis intermediates (Fig. 1a). Since *Toxoplasma* can ingest and digest host proteins to support its growth^10^, *Toxoplasma* may acquire heme through liberating it from the digestion of host’s hemoproteins. Collectively, both the parasite’s *de novo* heme production and/or heme acquisition from the host may contribute to parasite replication and infection. Interestingly, the *de novo* heme production is dispensable in the blood-stage infection of *Plasmodium falciparum*^11^, a malaria-causing agent, but is required for its liver-stage infection^12,13^, suggesting that intracellular pathogens could switch their metabolism for heme needs due to their access to exogenous heme availability.

To test whether all 8 genes residing within *Toxoplasma*’s *de novo* heme biosynthesis pathway are expressed in the acute infection stage, we endogenously inserted epitope tags at their C-termini (Fig. S1a). Immunoblotting revealed their active expression during acute toxoplasmosis (Fig. S1c). In addition, fluorescence localization experiments confirmed that *Toxoplasma* distributes its heme biosynthesis in the mitochondrion, cytoplasm, and apicoplast (Fig. 1b). Overall, our findings revealed that *Toxoplasma* maintains *de novo* heme biosynthetic components during acute infection.

Given the possibility that *Toxoplasma* could rely on its *de novo* heme production or scavenge heme or its intermediates from the host to support infection, we deleted 5-aminolevulinic acid (ALA) synthase (*TgALAS*, TGGT1_258690), the first enzyme in the pathway, in a NanoLuc luciferase-expressing wildtype (WT∷*NLuc*) *Toxoplasma*. Disruption of *TgALAS* will specifically block the parasite’s *de novo* heme biosynthesis but maintain the downstream pathway intact for possible utilization of host-derived heme intermediates (Fig. 1a). Generation of ∆*alas* in standard growth medium was unsuccessful until the medium was supplemented with 300 μM ALA, the product of TgALAS, suggesting that the *de novo* heme production is essential for parasite infection. To test this, we evaluated the replication of the resulting ∆*alas∷NLuc* mutant after it was starved in ALA-free medium for 144 h. The starved Δ*alas*∷*NLuc* exhibited severe replication defects compared to WT∷*NLuc* and Δ*alasALAS*∷*NLuc* (a *TgALAS* complementation strain), and also grew more slowly in the medium lacking ALA during replication relative to the starved Δ*alas*∷*NLuc* replicated in the ALA-containing medium (Fig. 2a). Additionally, the Δ*alas*∷*NLuc* parasites showed severe growth defects by using a luciferase-based growth assay (Fig. S5a). To further assess the viability of the Δ*alas*∷*NLuc* parasites in the absence of ALA, we monitored its growth over 10 days post-ALA starvation. The Δ*alas*∷*NLuc* mutant gradually slowed down its replication, but remained viable (Fig. 2b), suggesting that the parasites may incorporate a residual level of heme and/or heme synthesis intermediates from the host. To test the role of TgALAS in parasite acute virulence, we subcutaneously injected the Δ*alas*∷*NLuc* mutant along with WT∷*NLuc* and Δ*alasALAS*∷*NLuc* strains into CD-1 mice and did not observe mortality in the mice infected with the Δ*alas* strain, even when its inocula were 10^3^- and 10^4^-fold higher than that required for WT parasites to establish a lethal infection (Fig. 2c). As expected, the infections derived from WT∷*NLuc* and Δ*alasALAS*∷*NLuc* strains were lethal at 10-12 days post-infection (Fig. 2c). That surviving mice were infected was confirmed by seroconversion and their resistance to subsequent challenge with WT parasites.

**Fig. 2.**
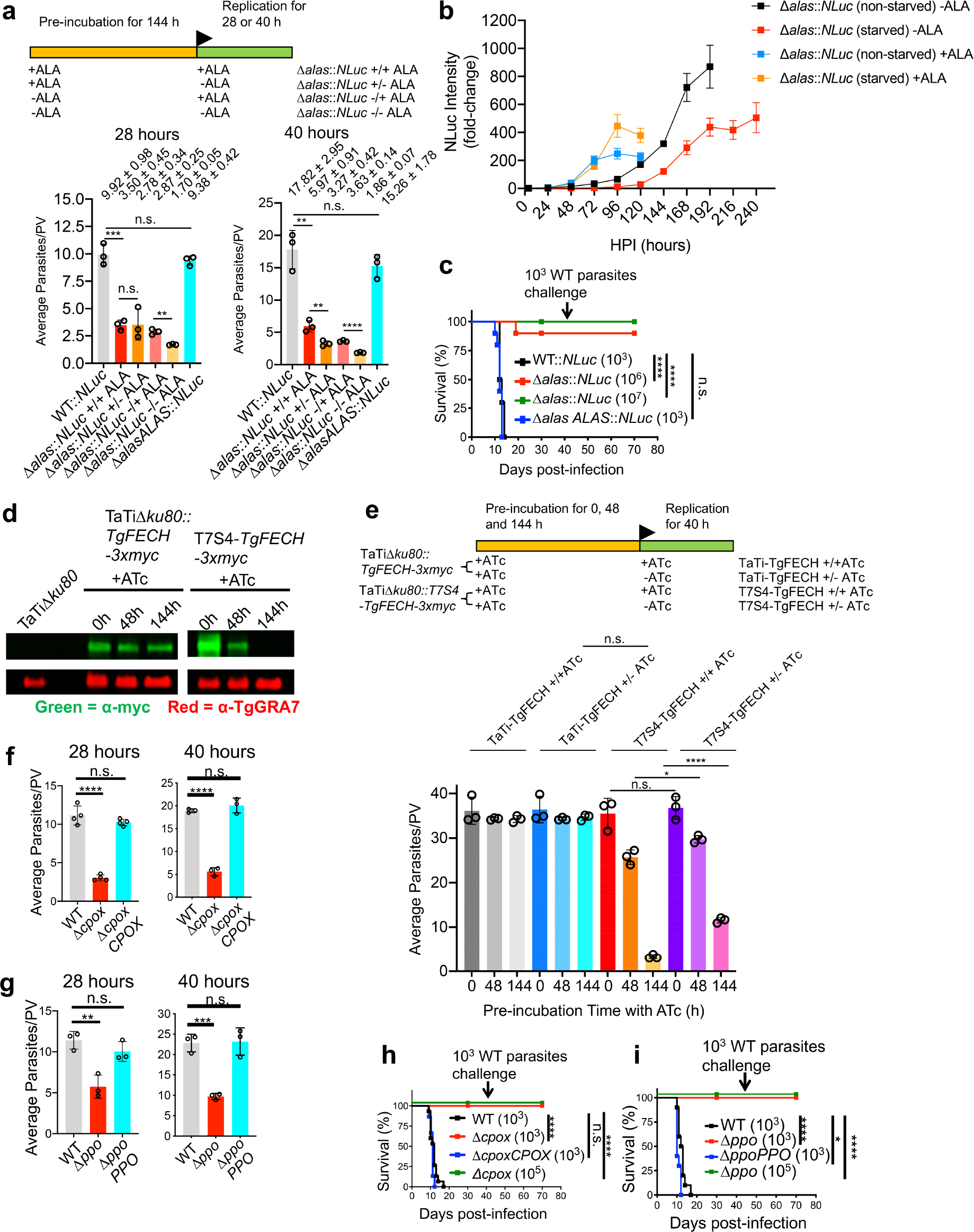
*Toxoplasma* parasites principally rely on their *de novo* heme biosynthesis for intracellular growth and pathogenesis. **a,** Replication comparison of the starved and non-starved Δ*alas*∷*NLuc* parasites in the media containing or lacking ALA. Data represent mean ± SD of n=3 biological replicates. **b,** A 10-day growth assay of the starved and non-starved Δ*alas*∷*NLuc* parasites in ALA-containing or deficient media. Error bars represent SEM of n=3 biological replicates with 3 technical replicates each. **c,** Acute virulence determination of *TgALAS*-deficient parasites in a murine model. Ten mice of equal numbers of males and females were used for each strain. **d,** Evaluation of repression efficiency of TgFECH by ATc treatment via immunoblotting. *TgFECH* was endogenously tagged with a 3xmyc tag at its C-terminus for recognition by immunoblotting. The lysates were also probed against TgGRA7 as loading control. **e,** Replication assessment of the *TgFECH* knockdown parasites. T7S4-TgFECH and its parental strains were pre-treated with ATc for the period described in the scheme before replication assay. Data represent mean ± SD of n=3 biological replicates. **f-g,** Replication assay of Δ*cpox* and Δ*ppo* parasites. Data represent mean ± SD of n=3-4 biological replicates. **h-i,** Acute virulence measurement of Δ*cpox* and Δ*ppo* parasites in a murine model. 5 male and 5 female mice were used for each strain. Statistical significance of the animal studies in **c, h,** and **i** was calculated using the Log-rank (Mantel-Cox) test. Statistical significance in the rest of the studies was calculated by two-tailed unpaired Student’s *t*-test. *, *p*<0.05; **, *p*<0.01; ***, *p*<0.001; ****, *p*<0.0001; n.s., not significant.

To further confirm that the heme biosynthesis pathway is crucial in *Toxoplasma*, we attempted to ablate the ferrochelatase (*TgFECH*, TGGT1_258650), which catalyzes the last step in heme production. We detected the correct integration of the drug resistance cassette into the *TgFECH* locus (Fig. S2d) but were unable to generate the straight Δ*fech* knockout, even upon supplementing the medium with 10 μM heme. Alternatively, we generated a *TgFECH* knockdown strain using a tetracycline-controlled TET-OFF system. To recognize the targeted *TgFECH*, we endogenously tagged its C-terminus with a 3xmyc epitope before replacing its cognate promoter with a T7S4 promoter, a hybrid of the promoter of *Toxoplasma SAG4* (surface antigen 4) gene and the ATc-responsive promoter (Fig. S3a). The expression of *TgFECH* in the resulting knockdown strain, named T7S4-TgFECH-3xmyc, was below the limit of detection after 144-h treatment of ATc (Fig. 2d), and the *TgFECH-*deficient mutant showed a drastic replication defect (Fig. 2e). Interestingly, if the *TgFECH* repression was halted during the replication assay by removing ATc, T7S4-TgFECH-3xmyc parasites significantly increased their replication rate compared to those grown in ATc-containing medium, validating the key role of the *de novo* heme biosynthesis in parasite intracellular growth (Fig. 2e). Additionally, we individually deleted coproporphyrinogen oxidase (*TgCPOX*, TGGT1_223020) and protoporphyrinogen oxidase (*TgPPO*, TGGT1_272490) within the pathway. The *TgCPOX*- and *TgPPO*-lacking mutants (Δ*cpox* and Δ*ppo*) displayed approximately 75% and 50% reduction in replication, respectively, compared to WT (Fig. 2f-g). A luciferase-based assay also confirmed that both knockout mutants showed severe growth defects compared to WT parasites (Fig. S5b-c). The deletion of *TgCPOX* or *TgPPO* also resulted in the complete loss of acute virulence in *Toxoplasma* (Fig. 2h-i). Given that both TgCPOX and TgPPO are oxidases, such oxidation reactions can occur even without enzymatic catalysis, albeit at lower reaction rates, which could explain why the growth defects of both knockouts were not as severe as that of *TgALAS* and *TgFECH*-deficient parasites. These findings are also consistent with *S. cerevisiae* wherein PPO is the only non-essential protein within its heme biosynthesis pathway^14^. In addition, *Toxoplasma* encodes an oxygen-independent coproporphyrinogen dehydrogenase (*TgCPDH*, TGGT1_288640), which may also help bypass the reaction catalyzed by TgCPOX to sustain partial *de novo* heme production in the parasites, albeit possibly at a lower level. Collectively, the systematic phenotypic characterization of a series of mutants lacking a functional *de novo* heme production in *Toxoplasma* established its crucial importance in parasite growth and pathogenesis. Although *Toxoplasma* may acquire heme or its intermediates from the host via a salvage pathway, it is not sufficient for supporting normal parasite growth and acute virulence.

To determine that the parasite’s *de novo* heme biosynthesis pathway is active, we complemented *S. cerevisiae* heme-deficient mutants lacking *ALAS*, *CPOX*, or *FECH* with the corresponding *Toxoplasma* ortholog genes. The resulting complementation strains had their heme auxotrophy phenotypes restored (Fig. 3a), whereas the strains complemented with the empty vector did not grow on heme-free media. These data, thus, established that TgALAS, TgCPOX, and TgFECH enzymes are functional. Additionally, the total heme abundance in heme biosynthetic gene knockout and knockdown parasites was quantified using a protoporphyrin IX-based fluorescence assay^14^. The total heme abundances in the Δ*cpox* and Δ*ppo* were reduced by approximately 75% and 50%, respectively (Fig. 3b). The heme level in Δ*alas* was nearly 50% relative to WT parasites when it was grown in medium supplemented with 300 μM ALA, but fell to approximately 10% when ALA was depleted (Fig. 3b). Similarly, the heme abundance in *TgFECH* knockdown strain was decreased to *ca.*15% compared to its parental strain when the mutant was treated with ATc for 144 h to repress *TgFECH* expression (Fig. 3b). Interestingly, the parasite replication rate was positively correlated to the total heme abundance in the parasites (Fig. 3b).

**Fig. 3.**
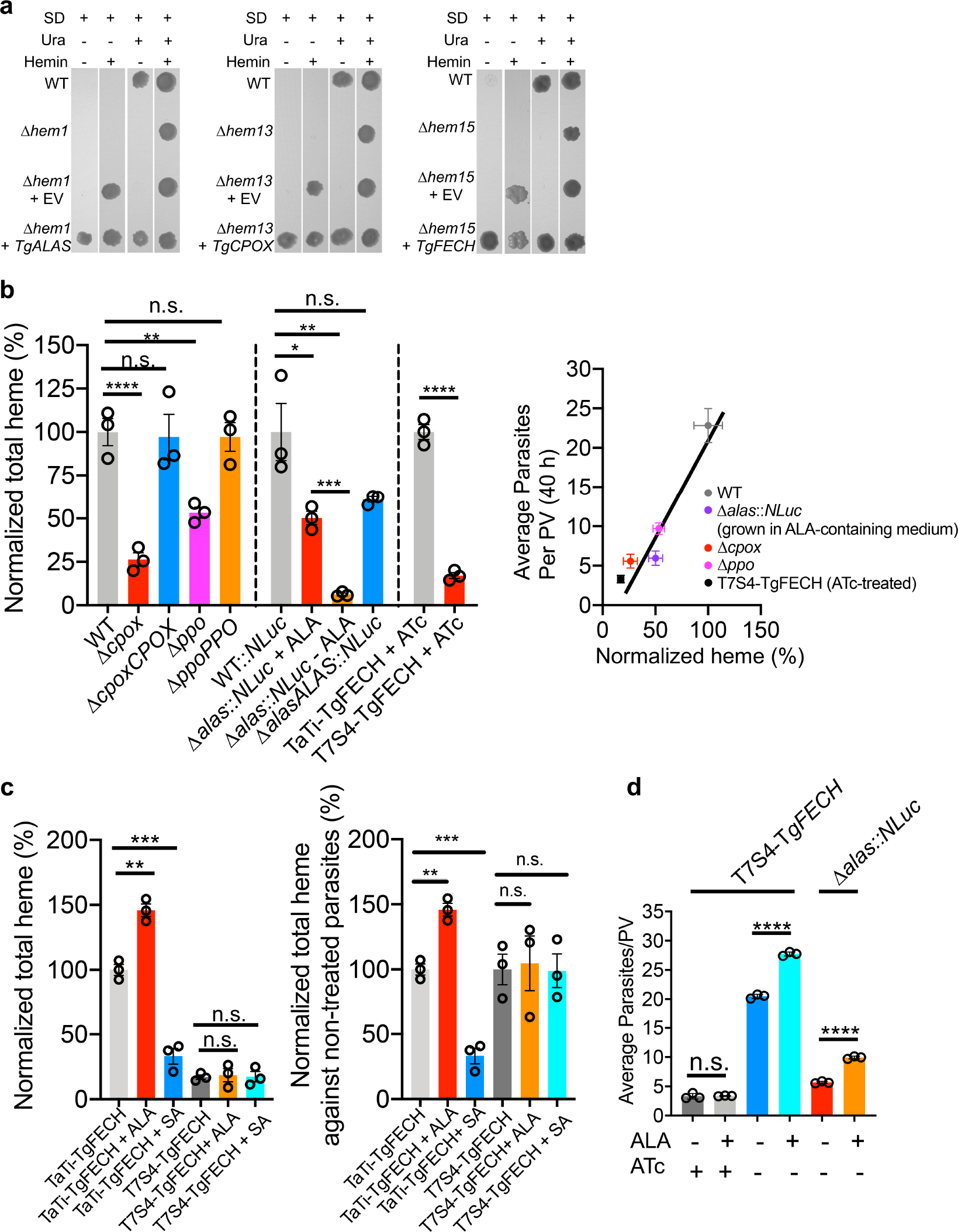
*Toxoplasma* harbors an active *de novo* heme biosynthetic pathway. **a,** Complementation of *Toxoplasma* orthologs of heme biosynthetic genes in the corresponding *S. cerevisiae* heme-deficient knockouts. EV, empty vector; SD, synthetic defined medium; Ura, uracil. **b,** Total heme quantification in heme-deficient parasites. A protoporphyrin IX-based fluorescence assay was used to quantify the total heme. The total heme levels in transgenic parasite strains were normalized against the corresponding parental strains. The average replication rates showed a positive correlation with the normalized heme abundances. Data represent mean ± SEM of n=3 biological replicates with 3 technical replicates each. **c,** Chemical interference in heme production in the parasites requires a functional heme biosynthetic pathway in *Toxoplasma*. The *TgFECH* knockdown parasites along with its parental strain were treated with ATc for 144 h before stimulation or repression of heme production by ALA or SA, respectively. Data represent mean ± SEM from n=3 biological replicates. **d,** Partial restoration of *TgFECH* expression in ATc-treated T7S4-TgFECH parasites helped them respond to the growth stimulation by ALA treatment. Data represent mean ± SD of n=3 biological replicates. Statistical significance in all of the studies listed in this figure was calculated by two-tailed unpaired Student’s *t*-test. *, *p*<0.05; **, *p*<0.01; ***, *p*<0.001; ****, *p*<0.0001; n.s., not significant.

We also evaluated whether the *de novo* heme production within *Toxoplasma* is responsive to chemical interference. Succinylacetone (SA), which inhibits *de novo* heme production by targeting TgPBGS activity, or ALA, which stimulates heme production, were used for chemical interrogation of the pathway (Fig. 1a). Initially, we determined that the concentration of SA that inhibited parasite growth by 50% (IC_50_) was 665.5 μM using a luciferase-based growth assay (Fig. S7). Next, we grew T7S4-TgFECH and its parental strain in the ATc-containing media for 144 h, followed by the addition of SA or ALA for an additional 48 h before heme quantification. The heme levels in parental parasites were increased ~40% by inclusion of ALA and reduced ~70% due to SA treatment (Fig. 3c), compared to DMSO (vehicle control)-treated parasites. However, ALA and SA did not alter heme levels in the ATc-treated T7S4-TgFECH strain (Fig. 3c). We also found that the inclusion of ALA in the medium improved parasite replication when the *de novo* heme biosynthesis pathway was partially restored in the T7S4-TgFECH strain (Fig. 3d). These findings suggest that the parasite’s *de novo* heme biosynthesis pathway actively responds to chemical stimuli for heme production and the fluctuation of heme abundance in host cells does not impact heme levels within the parasites. Taken together, these results indicated that *Toxoplasma* possesses an active heme biosynthesis pathway for their heme supply.

In plants, protoporphyrinogen oxidase (PPO) is involved in heme and chlorophyll production^15^. Chlorophyll is an essential pigment for photosynthesis in plants^16^. When PPO activity is inhibited in plants, the reactant of PPO, protoporphyrinogen IX, leaks into the cytoplasm and is spontaneously oxidized to protoporphyrin IX, which can absorb light to produce highly reactive singlet oxygen that destroys plant cell membranes^17^. Hence, PPO has been widely recognized as a target for herbicide development. A comparison of primary sequences of PPOs from mammals, plants, fungi, protozoans, and bacteria indicates that TgPPO is most closely related to plant orthologs (Fig. S8). Therefore, we evaluated 11 commercial herbicidal PPO inhibitors against WT *T. gondii* and identified that the IC_50_ values of 5 compounds ranged from ~130 to 650 μM for the inhibition of parasite growth as determined by a luciferase-based growth assay (Fig. S9).

Because oxadiazon was the most potent compound identified, and in light of its potential for additional medicinal chemistry optimization, we next evaluated two derivatives of the scaffold by modifying a structural homolog, oxadiargyl, by means of straightforward cycloaddition chemistry (Fig. 4a). Both derivatives had improved potency, with IC_50_ values approximately 15-25-fold lower than oxadiazon (Fig. 4b). In addition, we found that the Δ*ppo* parasites were less sensitive to these compounds than either WT or Δ*ppoPPO* strains (Fig. 4b). The differences in growth were not noted when WT or Δ*ppo* parasites were treated with pyrimethamine, a clinical antibiotic prescribed against acute toxoplasmosis by inhibiting folic acid synthesis (Fig. 4b). The Δ*ppoPPO* strain was less sensitive to pyrimethamine treatment because a pyrimethamine resistance cassette had been used to generate the *TgPPO* complementation plasmid. These findings suggest that these compounds were targeting TgPPO. Last, heme abundance in WT parasites treated with oxadiazon and its derivatives was reduced approximately 15-20% post-treatment (Fig. 4c), further supporting the notion that these chemicals targeted the heme biosynthesis pathway in the parasites. Notably, none of the compounds were toxic to human foreskin fibroblasts based on an AlarmarBlue-derived cell viability assay (Fig. S10).

**Fig. 4.**
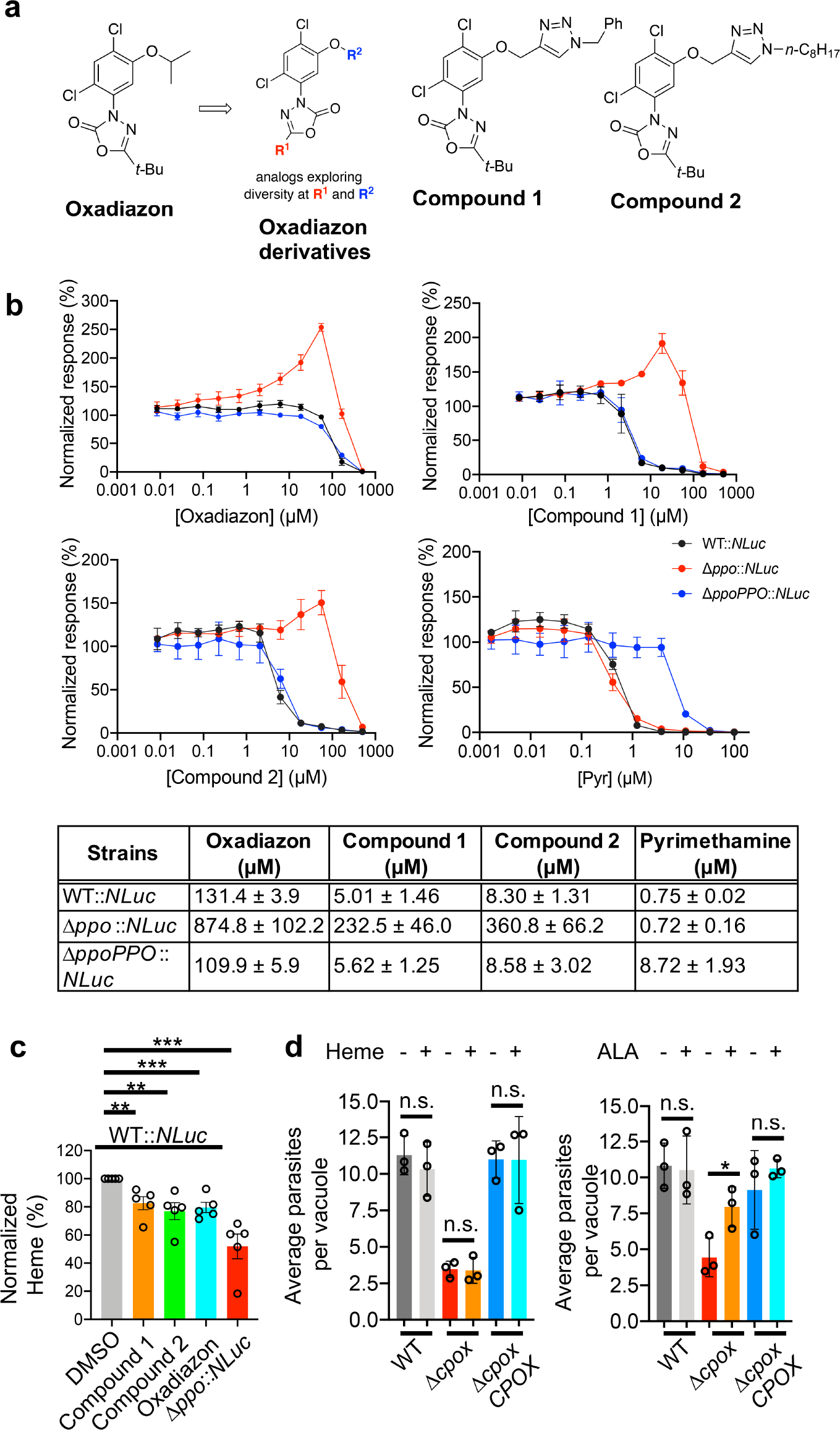
Chemical interrogation of the *Toxoplasma’s de novo* heme production by oxadiazon and its derivatives reduces the intracellular growth of the parasites. **a,** Chemical structure of oxadiazon and two derivatives. **b,** Efficacy determination of oxadiazon and its derivatives in the inhibition of WT *Toxoplasma* growth by using a luciferase-based growth assay. The Δ*ppo*∷*NLuc* and Δ*ppoPPO*∷*NLuc* were included for evaluating the target specificity of the inhibitors to TgPPO. Pyrimethamine, an antibiotic targeting a folic acid metabolism that is irrelevant to heme biosynthesis, was also included for assessing the specificity of these PPO inhibitors. Data shown in the table represent mean ± SEM of n=3 biological replicates with 3 technical replicates each. **c,** Heme levels were reduced in the parasites upon treatment with oxadiazon and its derivatives. Data shown here represent mean ± SEM of n=5 biological replicates. **d,** *Toxoplasma* was incapable of taking up extracellular heme to support its intracellular growth. Data represent mean ± SD of n=3 biological replicates. Statistical significance for the assays described in this figure was determined by two-tailed unpaired Student’s *t*-test. *, *p*<0.05; **, *p*<0.01; ***, *p*<0.001; n.s., not significant.

Given the unsuccessful attempts to delete *TgFECH* in the medium supplemented with heme, we speculate that *Toxoplasma* is unable to acquire adequate heme from the host to support its intracellular growth due to the lack of heme transporter. Supporting this, homology searches using several identified heme transporters as templates^18,19^, failed to identify candidate heme transporters in the *Toxoplasma* genome (ToxoDB database). We also used the Δ*cpox* cell line as a proxy to test whether *Toxoplasma* can incorporate extracellular heme to support intracellular replication. There was no significant difference in Δ*cpox* replication between standard growth medium and the medium supplemented with 10 μM heme (Fig. 4d). A Δ*cpox* replication assay between standard growth medium and the medium containing 300 μM ALA was included as a positive control (Fig. 4d) since *Toxoplasma* can incorporate extracellular ALA into its heme biosynthesis pathway as described above (Fig. 3c). A similar observation was seen in the intracellular growth of Δ*cpox* parasites between heme-depleted and heme-enriched media (Fig. S11). In contrast, humans can acquire dietary heme^18^ and mammalian cells can acquire extracellular heme via heme transporters^18,19^. Therefore, the host can overcome heme deficiency by heme supplementation during chemical inhibition of heme production, whereas the parasites cannot. These observations supported the concept that chemical interrogation of the parasite’s heme biosynthesis is a potential therapeutic strategy for controlling toxoplasmosis.

In conclusion, we revealed that *Toxoplasma* parasites harbor an active plant-like heme biosynthesis pathway and principally rely on this pathway to produce heme for their intracellular growth and acute virulence. In addition, the current antibiotics against *Toxoplasma* trigger strong side effects in some groups of patients and have limited efficacy on congenital toxoplasmosis. Thus, an urgent need for new therapeutics exists. Our findings shed light on the potential novel therapeutic targets within the parasite’s heme biosynthesis pathway for the clinical management of *Toxoplasma* infections.

## Materials and Methods

### Host cells and parasite culture

Human foreskin fibroblasts (HFFs) were obtained from the American Type Culture Collection (ATCC, catalog number: SCRC-1041). Tissue cultures were maintained at 37°C with 5% CO_2_ in D10 medium (Dulbecco’s Modified Eagle Medium, 4.5 g/L glucose, (VWR) supplemented with 10% Cosmic Calf serum (HycloneTM, GE Healthcare Life Sciences SH30087.03), 10 mM HEPES, additional 2 mM L-glutamine, and 10 mM Pen/Strep as instructed by the vendor’s manual. The HFF cells were tested for mycoplasma contamination every month. *Toxoplasma* strain RHΔ*ku80*Δ*hxg* and TaTiΔ*ku80*Δ*hxg* were obtained from the Carruthers Lab (Univ of Michigan) and the Striepen Lab (Univ of Pennsylvania), respectively, who originally created these *Toxoplasma* parental strains. All *Toxoplasma* strains were maintained *in vitro* by serial passage on HFFs. The Δ*alas*∷*NLuc* strain was kept in the D10 medium supplemented with 300 μM ALA.

### Generation of transgenic *T. gondii* strains

#### (1) Endogenously epitope-tagging the heme biosynthetic genes

The RHΔ*ku80*Δ*hxg* strain was used as the parental strain for endogenous gene tagging. The *TgALAS*, *TgALAD*, *TgCPOX*, *TgPPO*, and *TgFECH* were endogenously tagged with a 3xHA epitope at their C-termini by CRISPR-Cas9-mediated double crossover recombination as described previously^20^. The *TgPBGD*, *TgUROS*, and *TgUROD* were endogenously tagged with a 3xmyc tag C-terminally using similar methods as mentioned above. Briefly, the guide RNA selected to target the 3’-end of the coding sequence for individual genes was incorporated into a guide RNA expression construct as described previously^21^. This construct also encodes Cas9 protein that assists the guide RNA in generating a double-stranded break at the end of genes of interest. Fifty base pairs of homologous regions upstream and downstream stop codon were engineered into forward and reverse primers, respectively. By PCR, they flanked at the 5’- and 3’-ends of the epitope tag and a drug resistance cassette, respectively, to create a repair template. The guide RNA/Cas9 expression plasmids and the repair template were co-introduced into the parental strain by electroporation. Through double-crossover homologous recombination, the epitope tag and drug resistance cassette were incorporated after the last codon of individual genes (**Fig. S1a**). The resulting strains expressing epitope-tagged genes were drug selected and cloned out. The primers used for the epitope tagging of individual genes are listed in the **Supplementary Table 3**.

#### (2) Genetic ablation of the heme biosynthetic genes

A similar CRISPR-Cas9-based gene editing strategy was conducted for the deletion of *TgCPOX* and *TgPPO* in RHΔ*ku80*Δ*hxg* strain, and the ablation of *TgALAS* in RHΔ*ku80*∷*NLuc* strain generated in our previous study^20^. Similarly, the 50-bp homologous upstream and downstream DNA sequences of the start and stop codons of the corresponding gene, respectively, were incorporated into primers for amplification of the drug resistance cassette by PCR. In the final PCR product, the homologous regions located at both ends of the target gene facilitated the replacement of the gene of interest with a drug resistance cassette by double crossover recombination. For individual genes, the PCR product was mixed with the corresponding guide RNA used for the endogenous gene tagging and electroporated into RHΔ*ku80*Δ*hxg* or RHΔ*ku80*∷*NLuc* strains for knockout mutant generation. The correct clones were drug selected and cloned out as described previously^20^. PCR was used to verify the integration of the drug resistance cassette in the correct locus (**Fig. S2a-b**). The primers used for the deletion of individual genes and the verification of the knockout mutants are listed in the **Supplementary Table 3**.

#### (3) Complementation of the heme-deficient Toxoplasma knockouts

To restore the gene expression in the corresponding heme-deficient knockout, we PCR-amplified the coding sequences of the individual heme biosynthetic genes from the *Toxoplasma* cDNA library and the corresponding 1kb 5’- and 3’-UTR (untranslated regions) from the *Toxoplasma* genomic DNA. All three PCR products for individual genes were assembled into a plasmid vector carrying the pyrimethamine resistance cassette to create the corresponding complementation constructs. For both *TgPPO* and *TgALAS* complementation constructs, a 1kb DNA fragment localized around 6kb upstream region of the start codon of the *TgKU80* gene was included in the complementation construct to facilitate a single insertion of the complemented gene into the genome of the knockout strain to keep its expression similar to its endogenous expression level.

Approximately twenty micrograms of complementation plasmids were used for transfection. After stabilization by drug selection, the complementation parasites were cloned out. PCR was used to verify the integration of the introduced exogenous genes into the parasite’s genome. The primers used for the complementation of individual genes in the corresponding *Toxoplasma* knockout mutants and the verification of their integration are listed in the **Supplementary Table 3.**

#### (4) Generation of the ferrochelatase (TgFECH) knockdown strain using the tetracycline-controlled TET-OFF system

The TaTiΔ*ku80*Δ*hxg* strain carries the required genetic elements to suppress gene expression in response to the treatment of anhydrotetracycline (ATc)^22^, thereby being used as a parental strain to knock down *TgFECH* gene. To help recognize and evaluate the expression repression of *TgFECH*, we first endogenously tagged it with a 3xmyc epitope C-terminally as described above to create a TaTiΔ*ku80*∷*TgFECH-3xmyc* strain. A 1kb region upstream the start codon of *TgFECH* was recognized as the promoter region for a replacement with a tetracycline-responsive T7S4 promoter. To achieve this, the 50-bp regions upstream and downstream the *TgFECH* promoter region were incorporated at the 5’- and 3’-ends of the pyrimethamine resistance cassette and a tetracycline-responsive T7S4 promoter by PCR using a similar strategy as mentioned above. A single clone of TaTiΔ*ku80*∷*TgFECH-3xmyc* was transfected with the T7S4 promoter-encoding PCR product along with a guide RNA expression construct recognizing the 5’-end of *TgFECH* gene to create a T7S4-TgFECH-3xmyc strain. Please refer to **Fig. S3a** for the schematic description of the detailed steps. The epitope tag integration and the promoter replacement were confirmed by PCR (**Fig. S3b**). The primers used for the generation of the *TgFECH* knockdown strain and its verification are listed in the **Supplementary Table 3**.

#### (5) Generation of nanoLuc luciferase (NLuc)-expressing parasite strains

The *NLuc* expression plasmids were transfected into individual Δ*cpox* and Δ*ppo* parasite strains and the corresponding Δ*cpoxCPOX* and Δ*ppoPPO* complementation strains. The resulting *NLuc*-expressing strains were drug-selected and cloned out. Individual clones for each strain were screened by their luciferase activities to help select clones expressing similar levels of NLuc activities. These clones were maintained for downstream assays on intracellular growth and IC_50_ determination for PPO inhibitors.

### SDS-PAGE and immunoblotting

*Toxoplasma* parasites were grown and maintained in HFFs for a routine 2-day pass. Filter-purified parasites were resuspended in 1x SDS-PAGE sample buffer (40 mM Tris, pH 6.8, 1% SDS, 5% glycerol, 0.0003% bromophenol blue, 50 mM DTT) and boiled for 10 min prior to resolving on SDS-PAGE. Separated polypeptides were transferred to PVDF membranes via semi-dry protein transfer. For chemiluminescence-based detection, blots were blocked with 5% non-fat milk, incubated with primary antibody diluted in 1% non-fat milk in PBS, followed by probing with goat anti-mouse or anti-rabbit IgG antibodies conjugated with horseradish peroxidase as the secondary antibody diluted in 1% non-fat milk in PBS. Immunoblots were developed with SuperSignal WestPico chemiluminescent substrate (Thermo). For fluorescence-based detection, blots were blocked with 1.25% fish gelatin in 500 mM Tris, pH 7.4, 1.5 M NaCl, incubated with primary antibody diluted in wash buffer (0.1% Tween-20 in PBS), followed by probing with goat anti-mouse or anti-rabbit IgG antibodies conjugated with 680RD or 800CW dyes as the secondary antibody (LI-COR). The chemiluminescence and fluorescence signals were captured using the Azure C600 Imaging System.

### Immunofluorescence microscopy

Freshly lysed parasites were used to infect confluent HFF cells pre-seeded in an 8-well chamber slide for 1 h (pulse-invaded parasites) or 18-24 h (replicated parasites). Immunofluorescence was performed as described previously^20^. Images were viewed, digitally captured using a Leica CCD camera equipped with a Leica DMi8 inverted epifluorescence microscope, and processed with Leica LAS X software.

### PPIX-based total heme quantification

The parasites were grown in HFFs for 2 days in regular D10 medium or the medium supplemented with chemicals indicated in the text before harvest. Freshly lysed parasites were syringed, filter-purified, and washed in ice-cold PBS buffer. The parasites were finally resuspended in 400μl of ice-cold PBS buffer, and sonicated on ice using a Branson analog sonifier with a mini horn at the output intensity setting of 3 and duty% of 20% at a 10-sec interval for 4 times. A thirty-second rest was set between each sonication to avoid overheating. One hundred microliters of samples were mixed with 900 μl of 2 M oxalic acid in solid black Eppendorf tubes and boiled for 30 min. Similarly, each sample was duplicated in the same way without boiling, serving as a background. The heme solutions at 0, 5, 14.7, 44.3, 133, 400, and 1200 nM were included in the assay to produce a standard curve. The fluorescence was recorded at 400 nm excitation wavelength and 608 nm emission wavelength by BioTek H1 Hybrid plate reader. The fluorescence of non-boiled samples will be subtracted from that of boiled samples. The final fluorescence was normalized against the amount of parasite for comparison. The normalized heme abundance in WT parasites was set at 100% to calculate the relative heme abundance in other strains.

### Plaque assay of *T. gondii*

Freshly egressed parasites were filter-purified and resuspended in D10 medium. Two hundred parasites were inoculated into a 6-well plate seeded with confluent HFFs. The plate was incubated at 37°C with 5% CO_2_ for 7 days without disturbance. The developed plaques were stained with 0.002% crystal violet in 70% ethanol for 5 min and washed with water to enhance visualization of developed plaques. The plaques were scanned to capture the images of entire wells or observed under a microscope under 25x magnification by Leica DMi8 microscope. At least 50 individual plaques were photographed to measure their sizes. The plaque size was plotted for comparison.

### Replication assay of *T. gondii*

For the parasite replication assay, freshly lysed parasites were filter-purified and inoculated into individual wells of an 8-well chamber slide pre-seeded with HFF cells at approximately 1 × 10^5^ cells per well and incubated at 37°C with 5% CO_2_. Non-invaded parasites were washed off at 4 h post-infection. Invaded parasites were allowed to infect host cells for an additional 24 and 36 h before fixation. The infected host cells were stained with monoclonal anti-TgGRA7 (1:2,000) antibody and DAPI to help distinguish individual parasitophorous vacuoles (PVs) and parasite nuclei, respectively. Slides were subjected to standard immunofluorescence microscopy for imaging. One hundred PVs were enumerated for each strain and plotted as average parasites per PV for comparison.

### Bioluminescence-based growth assay

Individual strains were purified, resuspended in D10 medium, and inoculated into 96-well plate pre-seeded with confluent HFFs. Parasites were allowed to invade host cells for 4 h at 37°C with 5% CO_2_ before washing away non-invaded parasites, and the bioluminescence of invaded parasites at 4 h post-infection was measured for normalization. Invaded parasites incubated at 37°C with 5% CO_2_ for an additional 96 h, and their bioluminescence was measured every 24 h and normalized against the signal derived at 4 h for the calculation of the fold change of bioluminescence, which reflects parasite intracellular growth rate.

### Mouse studies

Six- to eight-week-old, outbred CD-1 mice were infected with WT or transgenic parasite strains diluted in PBS with inocula indicated in the text by subcutaneous injection. The infected mice were monitored daily for developing symptoms over 30 days. Mice that appeared moribund were humanely euthanized via CO_2_ overdose, in compliance with the protocol approved by Clemson University’s Institutional Animal Care and Use Committee (Animal Welfare Assurance A3737-01, protocol number AUP2016-012). The seroconversion of the surviving mice was tested by enzyme-linked immunosorbent assay (ELISA). The surviving mice were allowed to rest for 10 days, prior to subcutaneous challenge injection with 1000 WT parasites, and were kept for daily monitoring of survival for an additional 30 days. The mice survival curve was plotted for statistical significance calculation by using the Log-rank (Mantel-Cox) test.

### Yeast complementation assay

To test the functionality of the *Toxoplasma* orthologs of the heme biosynthetic proteins, we complemented these orthologs in the corresponding yeast knockouts. First, we PCR-amplified the G418 resistance cassette from the plasmid pFA6a-6xGLY-Myc-kanMX6 (Addgene, plasmid #:20769), flanked by 50-bp upstream and downstream regions of the start and stop codons of the corresponding heme biosynthetic gene, respectively. Approximately 5 μg of PCR products were transformed into the haploid yeast strain BY4741 (GE Healthcare Dharmacon Inc., catalog number: YSC 1048) to remove the coding sequences of heme biosynthetic genes by homologous recombination by using standard yeast transformation procedures (**Fig. S6a**). The transformants were selected on YPD plate containing 200 μg/ml of G418 and 15 μg/ml hemin. After two to three days of incubation, the correct yeast knockouts were identified by PCR (**Fig. S6b**). Next, we amplified the coding sequences of the corresponding *Toxoplasma* orthologs of individual heme biosynthetic genes from *Toxoplasma* cDNA library by PCR and drove its expression under the promoter of the yeast TEF1 gene in the pXP318 yeast expression construct to create a complementation construct. The yeast complementation construct was chemically introduced into the corresponding yeast knockout. The pXP318 encodes an uracil biosynthesis gene, while the WT yeast strain shows an uracil auxotrophy phenotype. Therefore, the successful transformants receiving pXP318 or pXP318-derived plasmids did not require the addition of uracil in the growth medium. The yeast knockout complemented with the corresponding *Toxoplasma* orthologs were also patched onto the plate lacking exogenously added heme to evaluate its heme autotrophy phenotype. The primers used for the yeast knockout generation and complementation of individual *Toxoplasma* orthologs are listed in the **Supplementary Table 3**.

### Synthesis of oxadiazon derivatives

All reagents were purchased from commercial sources and used without purification. ^1^H and ^13^C NMR spectra were collected on Bruker 300 MHz NMR spectrometers using CDCl_3_ as solvent. Chemical shifts are reported in parts per million (ppm). Spectra are referenced to residual solvent peaks. Infrared spectroscopy data were collected using an IR Affinity-1S instrument (with MIRacle 10 single reflection ATR accessory). Flash silica gel (40–63 μm) was used for column chromatography. The compounds were characterized by ^1^H and ^13^C NMR, ATR-FTIR, and HRMS. HRMS data were collected using an instrument equipped with electrospray ionization in positive mode (ESI+) and a Time Of Flight (TOF) detector (**Fig. S12 and S13**).

General Procedure for Oxadiargyl-triazole analogs^23^: Oxadiargyl (50 mg, 0.145 mmol, 1.0 equiv) was dissolved in DCM/H_2_O, 1:1 (0.5 mL) in a 20 mL round-bottomed flask equipped with a stir bar. The appropriate azide (0.175 mmol, 1.2 equiv) was added, followed by anhydrous CuSO_4_ (1 mg, 0.007 mmol, 0.05 equiv) and sodium ascorbate (4 mg, 0.02 mmol, 0.15 equiv). The resulting solution was then stirred vigorously at rt for 4 h. The reaction was diluted with DCM (5 mL) and water (5 mL). The organic layer was washed with saturated aq. brine, dried over anhydrous sodium sulfate, and concentrated by rotary evaporation to provide the crude product as a brown oil. The isolate was purified by silica gel flash column chromatography under the following gradient: hexanes (20 mL), ethyl acetate/hexanes (95:5; 20 mL), ethyl acetate/hexanes (90:10; 50 mL), ethyl acetate/hexanes (80:20; 50 mL), ethyl acetate/hexanes (70:30; 50 mL), ethyl acetate/hexanes (50:50; 50 mL) to afford an opaque oil (62–87% isolated yield).

#### 3-(5-((1-benzyl-*1H*-1,2,3-triazol-4-yl)methoxy)-2,4-dichlorophenyl)-5-(*tert*-butyl)-1,3,4-oxadiazol-2(*3H*)-one (**Compound 1**)

Compound 1 was obtained in 62% yield using benzyl azide^24^ (23 mg, 0.18 mmol, 1.2 equiv) as the azide reagent.

Opaque oil; Yield: 62% (69 mg); R_*f*_ 0.54 (1:1 ethyl acetate/hexanes, UV); IR: (film) □ = 2970, 2924, 2856, 2094, 1782, 1489, 1246, 752, 717 cm^−1^; ^1^H NMR: (300 MHz, CDCl_3_) δ 1.39 (s, 9H), δ 5.29 (s, 2H), δ 5.55 (s, 2H), δ 7.24 (s, 1H), δ 7.30 (m, 2H), δ = 7.38 (m, 2H), δ 7.52 (s, 1H), δ 7.61 (s, 1H); ^13^C{^1^H} NMR: (75 MHz, CDCl_3_) δ = 27.0, 32.9, 54.3, 63.7, 113.8, 123.0, 123.9, 125.2, 128.1, 128.9, 129.2, 131.4, 134.3, 142.6, 152.0, 152.9, 163.6; HRMS (ESI-TOF): Calcd. for C_22_H_22_Cl_2_N_5_O_3_, [M+H]^+^ 474.1100 found m/z 474.1099.

#### 5-(*tert*-butyl)-3-(2,4-dichloro-5-((1-octyl-*1H*-1,2,3-triazol-4-yl)methoxy)phenyl)-1,3,4-oxadiazol-2(*3H*)-one (**Compound 2**)

Compound 2 was obtained in 87% yield using octyl azide^25^ (24 mg, 0.18 mmol, 1.2 equiv) as the azide reagent.

Opaque oil; Yield 87% (72 mg); R_*f*_ 0.50 (1:1 ethyl acetate/hexanes, UV); IR: (film) □ = 2951, 2924, 2854, 1786, 1479, 1246, 1126, 1041, 733 cm^−1^; ^1^H NMR: (300 MHz, CDCl_3_) δ = 0.87 (t, 3H, *J* = 7 Hz), 1.26 (m, 12H), 1.38 (s, 9H), 1.92 (p, 2H, *J* = 7 Hz), 4.36 (t, 2H, *J* = 7 Hz), 7.26 (s, 1H), 7.52 (s, 1H), 7.68 (s, 1H); ^13^C{^1^H} NMR: (75 MHz, CDCl_3_) δ = 27.0, 32.9, 54.3, 63.7, 113.8, 122.9, 123.9, 125.1, 131.4, 142.6, 152.0, 152.9, 163.6; HRMS (ESI+-TOF): Calcd for C_23_H_31_Cl_2_N_5_O_3_, [M+H]^+^ 496.1882 Found m/z 496.1878.

### *In vitro* measurement of IC_50_s for chemical inhibitors

General procedures: Individual parasite strains expressing NanoLuc luciferase were used to infect confluent HFFs pre-seeded into 96-well solid white plate with inocula of 1000 tachyzoites per well for WT∷*NLuc* and Δ*ppoPPO*∷*NLuc* strains and 3000 tachyzoites per well for Δ*ppo*∷*NLuc* strain. Higher inoculum of Δ*ppo*∷*NLuc* were adjusted in the assay to help accurately quantify fold change of luciferase activity due to its slow growth. Purified parasites were resuspended in phenol red free D10 medium, and 150 μl of parasite resuspension for individual strains were inoculated each well. Parasites were initially incubated at 37°C with 5% CO_2_ for 4 h to facilitate host invasion. Next, the media were gently aspirated and switched to the phenol red free D10 media containing individual chemical inhibitors at the serial concentrations listed below. A control medium without an inhibitor was included for luciferase activity normalization. After 2- to 4-day incubation with chemical inhibitors, the media were gently aspirated to avoid disturbance of monolayer HFFs, changed to the 12.5 μM Coelenterazine h in lysis buffer (100 mM 4-Morpholineethanesulfonic acid (MES) pH 6.0, 1 mM trans-1,2-Diaminocyclohexane-N,N,N′,N′-tetraacetic acid (CDTA), 0.5% (v/v) Tergitol, 0.05% (v/v) Mazu DF 204, 150 mM KCl, 1 mM DTT, and 35 mM thiourea), and incubated at room temperature for 10 min to fully lyse cells prior to luciferase activity quantification. The signals under individual concentrations of inhibitors were recorded by BioTek plate reader and normalized against the luciferase activities derived from the infected cells incubated in the plain medium. The normalized luciferase activities were plotted against the concentrations of inhibitors to calculate the IC_50_ values for individual inhibitors by the built-in “Dose-response-inhibition” program in GraphPad Prism software (8^th^ version).

#### Measurement of IC_50_s for the PPO inhibitors

The serial concentrations of PPO inhibitors were 500.0, 125.0, 31.3, 7.81, 1.95, 0.488, 0.122, 3.05×10^−2^, and 7.63×10^−3^ μM. The efficacies of these inhibitors were only tested on WT∷*NLuc* parasites after 4 days of growth.

#### Measurement of IC_50_s for Oxadiazon and its derivatives

The serial concentrations of the chemicals were 500.0, 166.7, 55.6, 18.5, 6.17, 2.06, 6.8×10^−1^, 2.3×10^−1^, 7.6×10^−2^, 2.5×10^−2^, and 8.5×10^−3^ μM. The efficacies of these inhibitors were tested on WT∷*NLuc*, Δ*ppo*∷*NLuc*, and Δ*ppoPPO*∷*NLuc* parasites after 4 days of growth.

#### Measurement of IC_50_ for pyrimethamine

The serial concentrations of pyrimethamine included in the media were 100.0, 33.3. 11.1, 3.70, 1.23, 0.41, 0.14. 0.045, 0.015, 5.1×10^−3^, and 1.7×10^−3^ μM. The efficacy of pyrimethamine was tested on WT∷*NLuc*, Δ*ppo*∷*NLuc*, and Δ*ppoPPO*∷*NLuc* parasites after 4 days of growth.

#### Measurement of IC_50_ for succinylacetone

The serial concentrations of pyrimethamine included in the media were 100.0, 33.3. 11.1, 3.70, 1.23, 0.41, 0.14. 0.045, 0.015, 5.1×10^−3^, and 1.7×10^−3^ μM. The IC_50_ value of succinylacetone was determined on its inhibition on 2-day growth of WT∷*NLuc* parasites.

### Resazurin-based cell viability assay

The HFFs were seeded in 96-well plates and grown in regular D10 medium. After the host cells became confluent, they were incubated in the media containing the tested PPO inhibitors in the serial concentrations at 500.0, 166.7, 55.6, 18.5, 6.17, 2.06, 6.8×10^−1^, 2.3×10^−1^, 7.6×10^−2^, 2.5×10^−2^, and 8.5×10^−3^ μM, or the media containing Triton X-100 at 1070.0, 356.7, 118.9, 39.6, 13.2, 4.40, 1.47, 0.49, 0.16, 0.054, and 0.018 μg/ml. The host cells grown in regular D10 medium were included as the control for normalization. The resazurin was diluted at 0.004% in regular D10 medium and incubated with host cells at 37°C with 5% CO_2_ for 4 h before absorbance measurement at 570 and 600 nm by BioTek H1 Hybrid plate reader. A resazurin-containing medium control without incubation with host cells was included to calculate the absorbance ratio of oxidized resazurin at 570 nm vs. 600 nm, termed OD(blank)_570/600_. The normalized cell viability was calculated using the following equation: [OD(inhibitor)_570_ − OD(inhibitor)_600_ × OD(blank)_570/600_]/ [OD(medium)_570_ − OD(medium)_600_ × OD(blank)_570/600_] *100%.

### Phylogenetic tree construction

Polypeptide sequences of 17 PPO orthologs from animals, plants, bacteria, protozoa, and fungi were retrieved from the Uniport database (www.uniprot.org). The sequences were aligned using the CLUSTAL MUSCLE (MUltiple Sequence Comparison by Log-Expectation) alignment tools^26,27^. The resulting sequence alignment was used to construct a phylogenetic tree by using the Neighbor-Joining tree analysis^28^ built in the Geneious software. Bootstrap values based on 10,000 replicates are shown. The sequences used for MUSCLE alignment were listed in **Fig. S8**.

### Statistical significance calculation

The statistical significance calculation in this study was performed using GraphPad Prism software (8^th^ version). The methods for individual assays were indicated in the figure captions.

## Data availability

The data that support the findings of this study are available from the corresponding authors upon request.

## Acknowledgements

We thank our colleagues, Drs. Vern Carruthers, David Sibley, and Peter Bradley for sharing key reagents for this work. We also want to thank Drs. Vern Carruthers, James Morris, and Meredith Morris for critically reading this manuscript before submission. This work was supported by the Clemson Startup fund (to Z.D.), Knights Templar Eye Foundation Pediatric Ophthalmology Career-Starter Research Grant (to Z.D.), a pilot grant of an NIH COBRE grant P20GM109094 (to Z.D.), NIH R01AI111962 and R01AI056312 (to I.H.). The funders had no role in study design, data collection and analysis, decision to publish, or preparation of the manuscript.

## Author contributions

Z.D. designed this study. A.B., K.F., M.K., C.D, K.C.R., and Z.D. executed experiments. A.B., K.F., and Z.D. performed endogenous gene tagging and determined their subcellular locations by immunofluorescence microscopy and their expression in acute *Toxoplasma* infection via immunoblotting. A.B., K.F., C.D., and Z.D generated complete knockout or knockdown parasite strains and the corresponding complementation strains if needed. A.B., K.F., M.K., C.D., and Z.D. measured the replication, growth, and acute virulence for the heme-deficient parasites. A.B., K.F., and C.D. conducted plaque assays for the heme-deficient parasites. Z.D. generated the heme-deficient yeast strains and trans-complemented them with the corresponding wildtype *Toxoplasma* orthologs. A.B., K.F., M.K., and Z.D quantified heme abundances in the heme-deficient and drug-treated parasites. K.F. and Z.D. evaluated the inhibition efficacies of the PPO-targeting herbicides and their derivatives. K.C.R. and D.C.W. designed the strategy for chemical synthesis of oxadiazon derivatives and conducted the chemical modification. I.Q. provided analytic methods for heme quantification and technical support to this study. Z.D. wrote the original draft. A.B., K.F., M.K., C.D., K.C.R., D.C.W., I.Q., and Z.D. reviewed and revised the manuscript.

## Materials & Correspondence

Please send inquiries to Zhicheng Dou

