## Supplementary text 1 for "*Toxoplasma gondii* requires its plant-like heme biosynthesis pathway for infection"

### SUPPLEMENTAL TEXT 1

#### 1. TgPBGD is an apicoplast-localizing enzyme

A previous study of overexpression of TgPBGD revealed its subcellular location in the cytoplasm of the parasites[4]. By endogenous gene tagging, we determined its subcellular location in the apicoplast (**Fig. 1b and S1b**). Therefore, TgCPOX is the only cytosolic enzyme in the parasite's heme biosynthetic pathway.

#### 2. Generation of the heme-deficient *Toxoplasma* strains

We used the CRISPR-Cas9-based genome editing tools to ablate the entire coding region, including introns and exons of 3 heme biosynthetic genes: *TgALAS*, *TgCPOX*, and *TgPPO*. By PCR, we confirmed the removal of the endogenous genes and the integration of the drug resistance cassette (**Fig. S2b**). Additionally, the total RNA from the knockouts, their parental strains, and the corresponding complementation strains were purified and tested by RT-PCR to confirm the loss of their messenger RNAs in the knockouts and the reversal expression in the complementation strains (**Fig. S2c**). We used a similar strategy to remove *TgFECH* coding sequence in the parasites. After two rounds of drug selection, the integration of the bleomycin cassette into the *TgFECH* locus was detected by PCR (**Fig. S2d**). However, we were unable to isolate the correct knockout clone from the population, suggesting that the *TgFECH* location in the genome is accessible for homologous recombination; however, *TgFECH* is required for parasite growth.

#### 3. The heme-deficient *Toxoplasma* parasites form small plaques.

The heme-deficient parasites were used to infect confluent HFFs in 6-well plates. The plates were incubated at 37ºC with 5% $CO_2$ for 7 days before plaque development was observed by crystal violet staining. The sizes of the plaques formed by Δ*cpox* and Δ*ppo* were reduced by ~90% and ~70%, respectively, compared to WT parasites (**Fig. S4a-b**). For the *TgALAS*-deficient parasites, we grew the parasites in media containing or lacking ALA. The addition of extracellular ALA improved parasite growth. In the absence of ALA, the plaques formed by Δ*alas*::*NLuc* parasites showed small dark regions. In the presence and absence of ALA, the plaque areas of Δ*alas*::*NLuc* parasites were reduced by ~98% and 87%, respectively, relative to WT parasites. Given that the replication rates of Δ*cpox* and Δ*ppo* were only reduced by ~75% and 50%, respectively, compared to WT parasites (**Fig. S4c**), these findings suggest that the heme-deficient parasites may display defects in other steps of the parasite's lytic cycle, such as invasion and egress.

**SUPPLEMENTARY TEXT 2**

**1. Primers used in *Toxoplasma* knockout generation in Fig. S2.**
For Δ*alas*::*NLuc*, P1, CUP41; P2, P414; P3, P415; P4, CUP42; P5, CUP43; P6, CUP44.

For Δ*cpox*, P1, P467; P2, P414; P3, P415; P4, P468; P5, CUP223; P6, CUP224.

For Δ*ppo*, P1, CUP241; P2, P414; P3, P415; P4, CUP242; P5, CUP239; P6, CUP240.

For Δ*fech*, P1, CUP363; P2, P414; P3, P415; P4, CUP364.

**2. Primers used in yeast knockout generation in Fig. S6.**
For Δ*hem1*, P1, CUP554; P2, CUP302; P3, CUP303; P4, CUP555; P5, CUP561; P6, CUP560.

For Δ*hem13*, P1, CUP300; P2, CUP302; P3, CUP303; P4, CUP301; P5, CUP727; P6, CUP728.

For Δ*hem15*, P1, CUP705; P2, CUP302; P3, CUP303; P4, CUP706; P5, CUP703; P6, CUP704.

The CUP292 and CUP293 primers were used to test the introduction of the *Toxoplasma* orthologs in the trans-genera complementation strains.
