## Supplementary figures for "*Toxoplasma gondii* requires its plant-like heme biosynthesis pathway for infection"

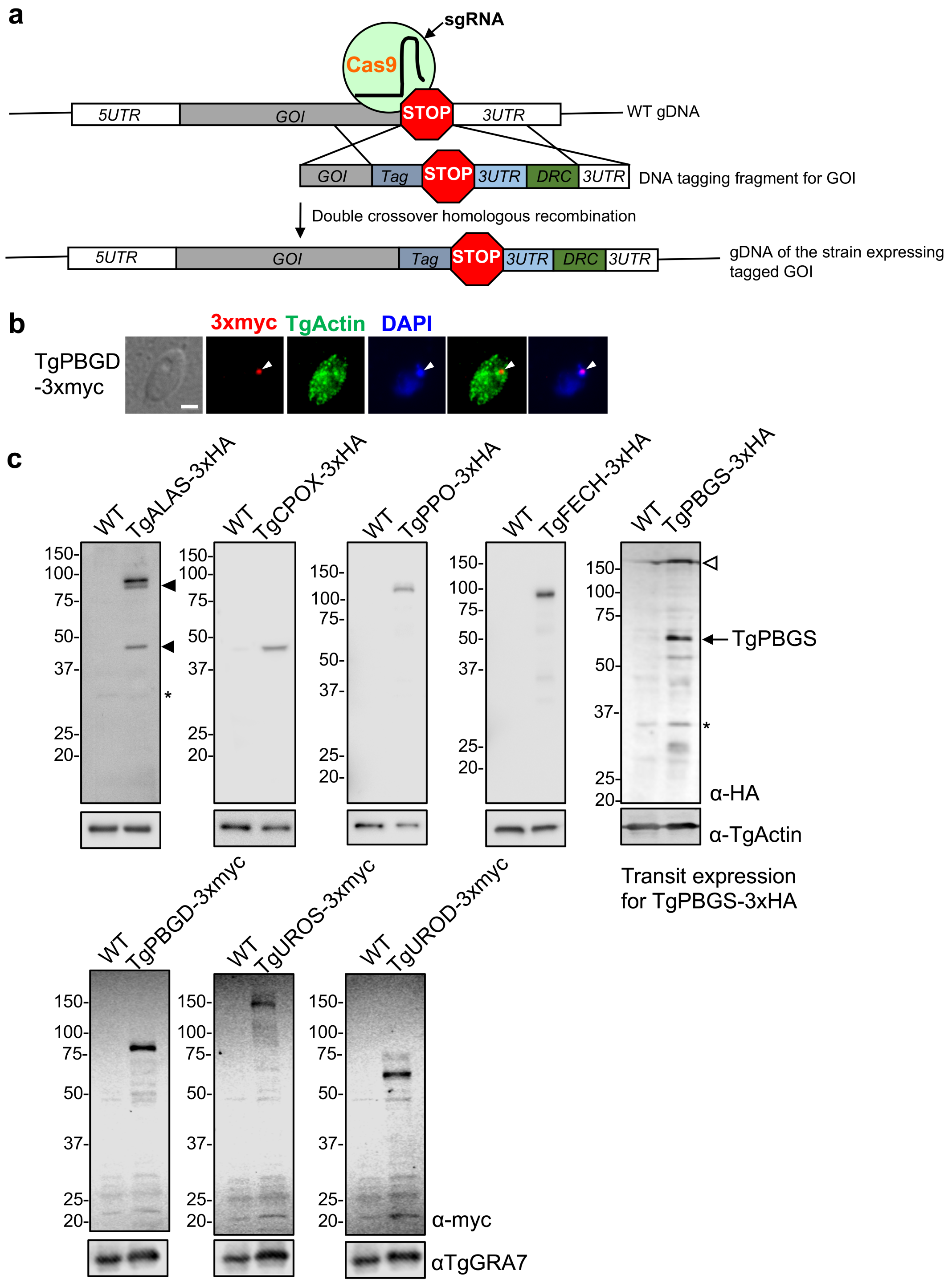


**Fig. S1 | Endogenous epitope tagging of the *Toxoplasma* heme biosynthetic genes. a,** Schematic illustration of the endogenous epitope tagging in *Toxoplasma*. In brief, a 3xHA or 3xmyc tag was fused at the C-termini of the genes of interest by CRISPR-Cas9-based cloning strategy. **b,** TgPBGD was localized to the apicoplast, instead of the cytoplasm. TgActin was used as a cytoplasm marker. Bar = 2 µm. **c,** Immunoblotting analysis was used to confirm the expression of the epitope-tagged genes. The bands labeled with asterisks were derived from non-specific binding to antibodies. For TgALAS-3xHA, two protein fragments denoted by the filled arrowheads, a ~80 kDa band close to the major band and a band migrating ~40 kDa, were the truncated forms of TgALAS. The *TgPBGS* gene only can be endogenously epitope-tagged in a transient manner. The lysate was purified from the parasites lysed immediately after transfection with the guide RNA expression construct and a TgPBGS-3xHA tagging DNA fragment. The guide RNA expression construct also expressed the 3xHA-tagged Cas9 proteins. Based on the predicted molecular weight, the band denoted by an unfilled arrowhead was derived from 3xHA-tagged Cas9. GOI, gene of interest; DRC, drug resistance cassette.

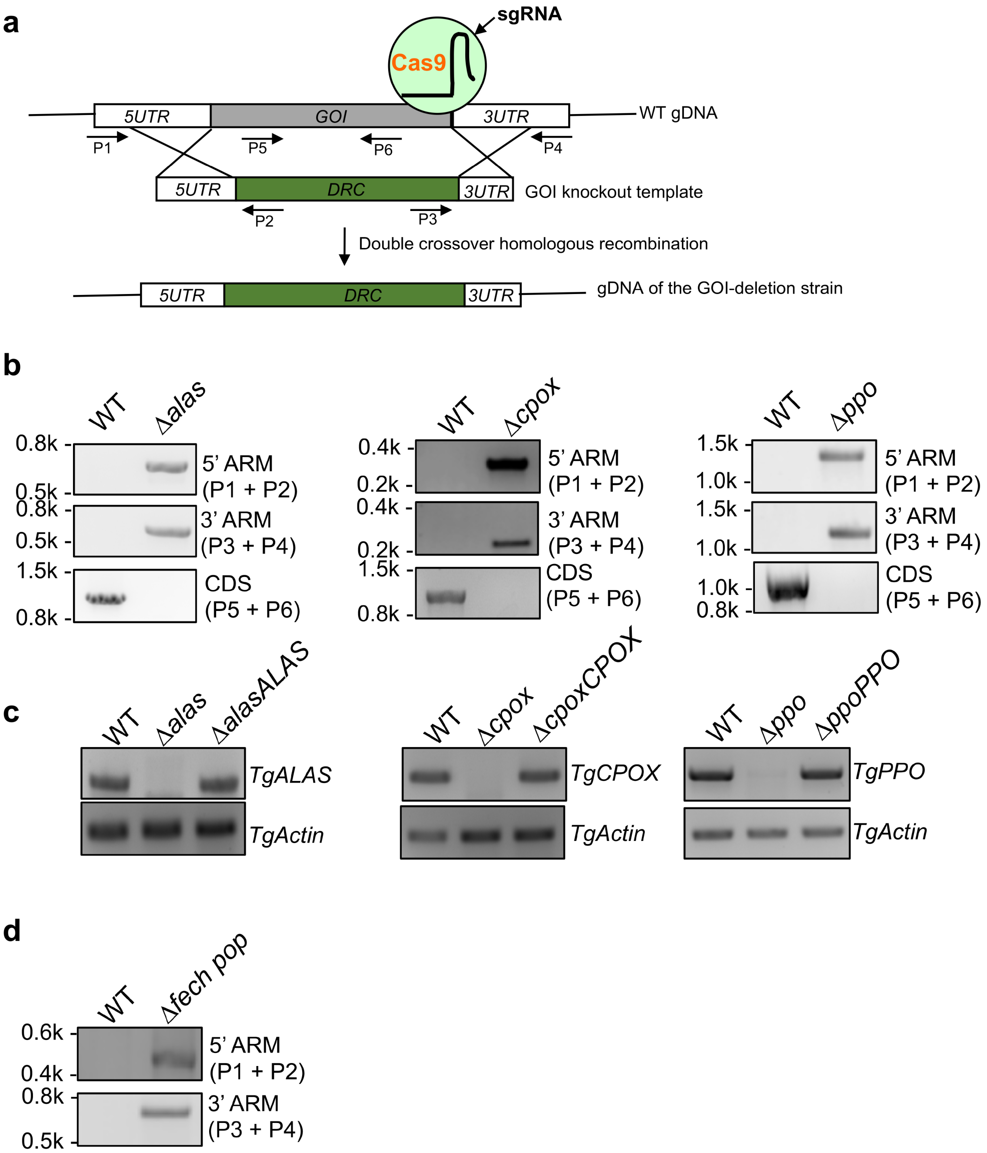


**Fig. S2 | Genetic deletion of the *Toxoplasma* heme biosynthetic genes. a,** Schematic illustration of a general CRISPR-Cas9-based strategy for gene deletion in *Toxoplasma*. **b,** PCR confirmation of gene ablation. The genomic locations of the primers used in PCR amplification were indicated in the scheme. **c,** The loss of messenger RNA of the genes of interest was confirmed by reverse-transcription PCR (RT-PCR). **d,** The correct integration of the drug resistance cassette into the *TgFECH* locus was detected by PCR during gene deletion. However, the knockout parasites cannot be cloned probably due to its non-viability. GOI, gene of interest; DRC, drug resistance cassette.

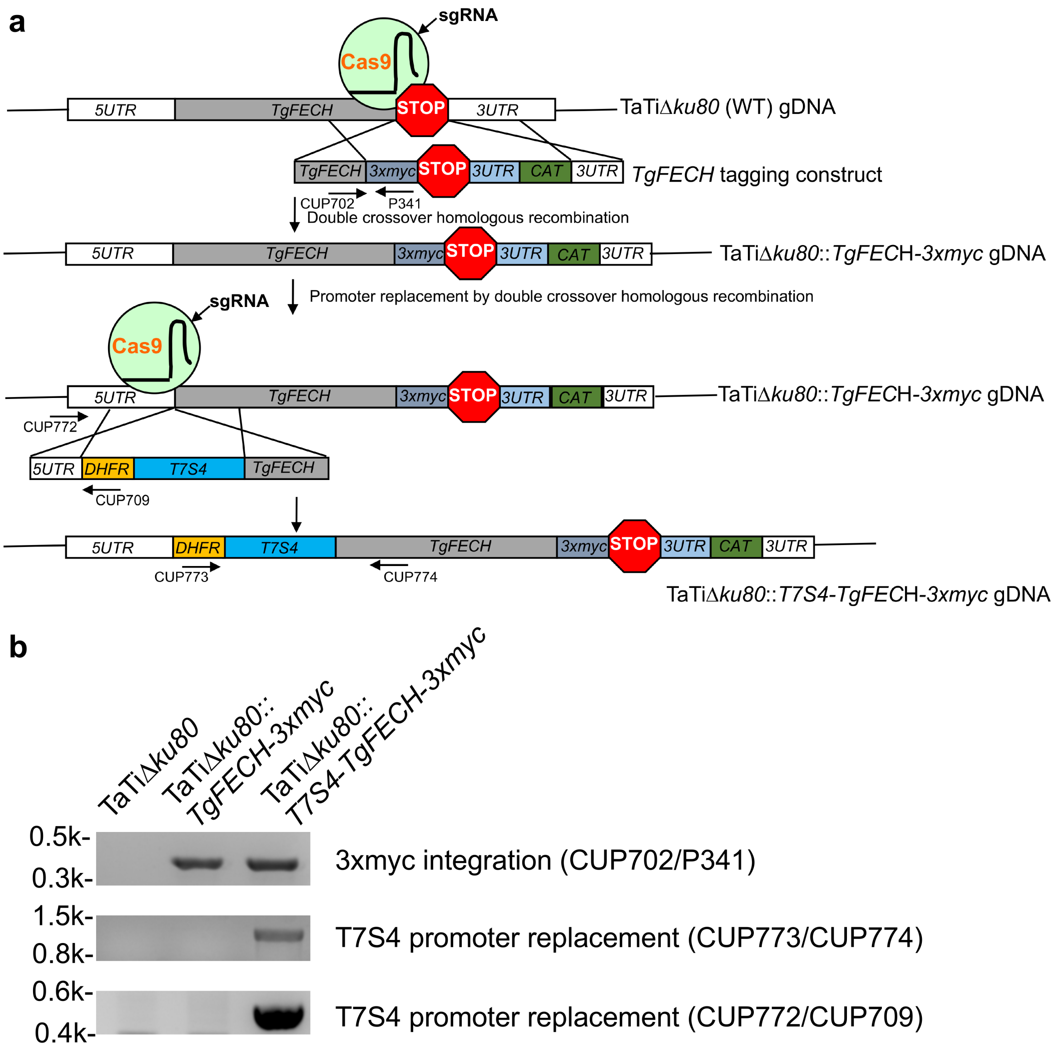


**Fig. S3 | *TgFECH* expression was regulated by a tetracycline-inducible TET-OFF system. a,** Graphic description of gene epitope tagging and promoter swapping for *TgFECH* gene. **b,** PCR verification of the integration of the 3xmyc tag and TET-OFF promoter into the *TgFECH* locus. Primers used in this study were indicated in the scheme.

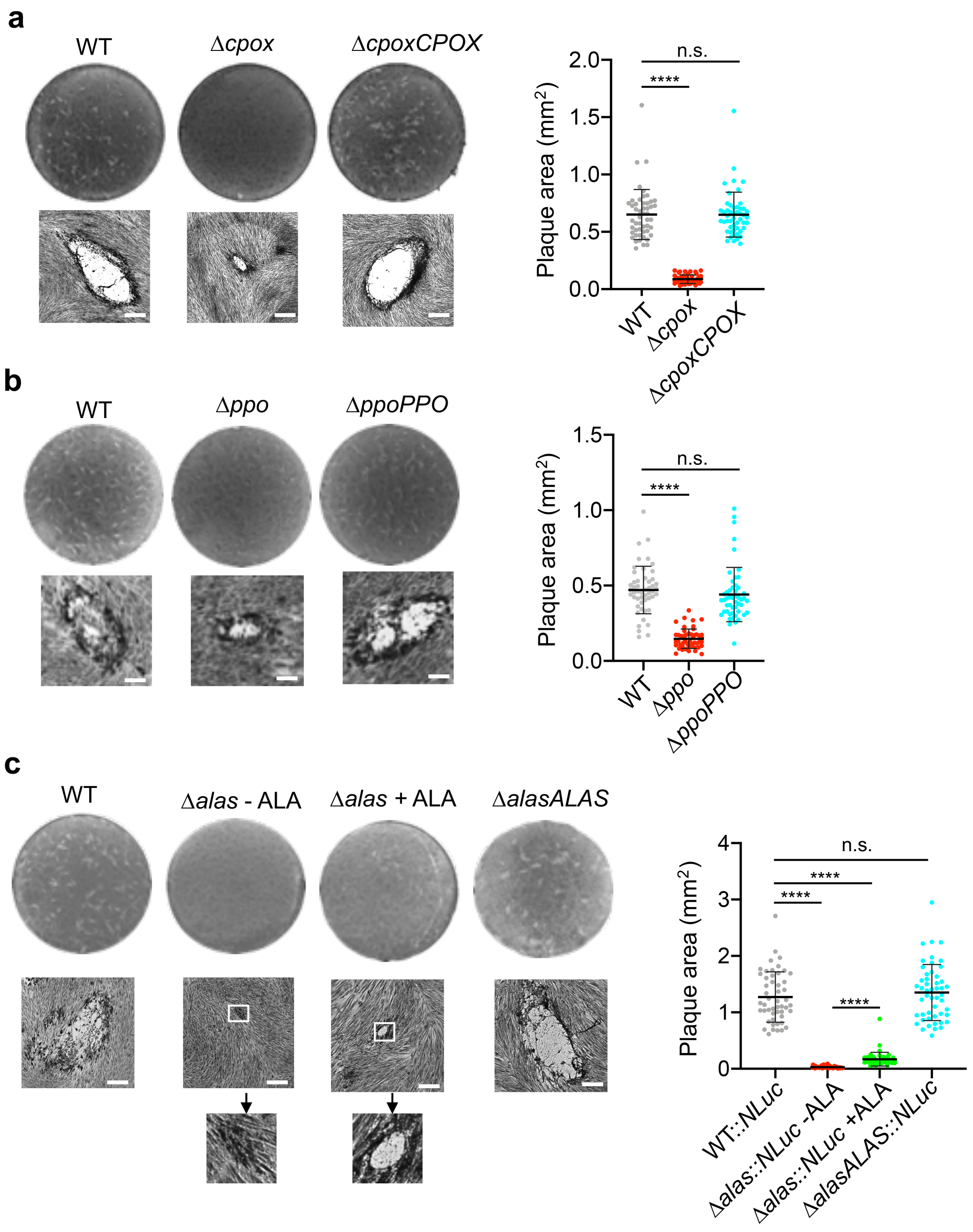


**Fig. S4 | Plaque assay of the *Toxoplasma* heme-deficient parasites. a,** The Δ*cpox* displayed smaller plaques than WT and Δ*cpoxCPOX* parasites. The plaques were allowed to develop in confluent HFFs for 7 days without disturbance before staining with crystal violet. Fifty plaques from 3 independent assays were measured under phase contrast light microscope to compare their sizes. Bar = 500 μm. Data represent mean ± SD. **b-c,** Plaque assays for *TgPPO*- and *TgALAS*-deficient parasites. Statistical significance was calculated by unpaired Student's *t*-test. ****, $p<0.0001$; n.s., not significant.

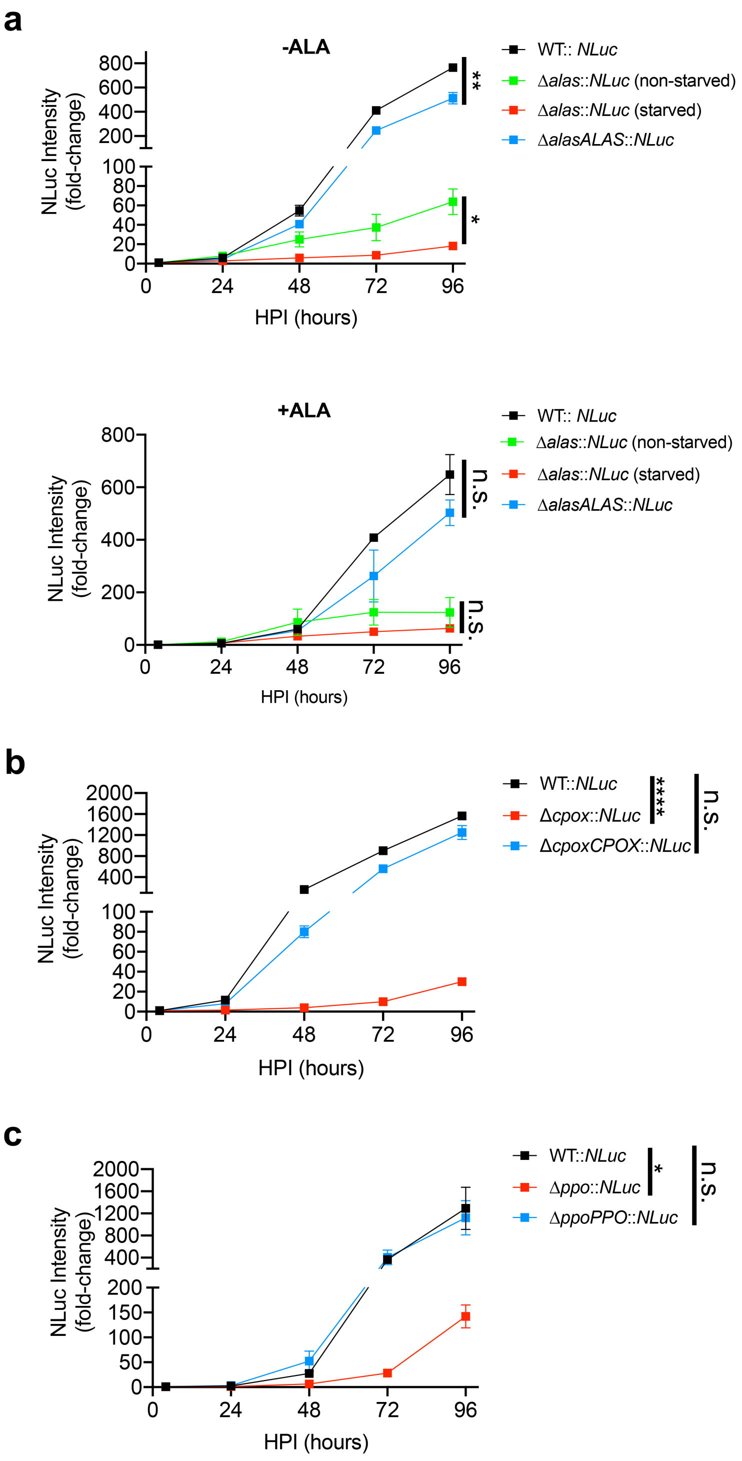


**Fig. S5 | Intracellular growth determination of heme-deficient parasites using a luciferase-based assay. a-c,** The heme-deficient parasites were grown in confluent HFFs and their luciferase activities were measured every 24 h up to 96 h. The luciferase activities at 4 h post-infection were also determined for normalization. Error bars represent SEM. The assays were repeated in triplicate. Statistical significance was determined by unpaired Student's *t*-test. *, $p$<0.05; **, $p$<0.01; ****, $p$<0.0001; n.s., not significant.

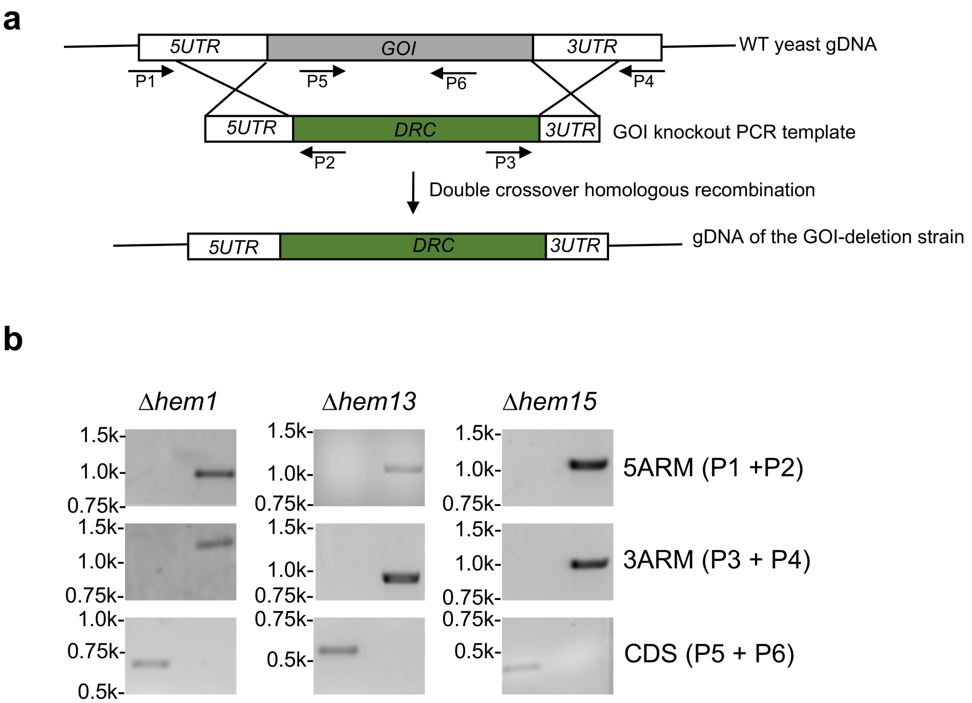


**Fig. S6 | Generation of heme-deficient yeast strains. a,** Schematic illustration of the gene deletion strategy. **b,** PCR was used to verify the loss of the heme biosynthetic genes in yeast. Primers used in the study were labeled in the scheme. GOI, gene of interest; DRC, drug resistance cassette.

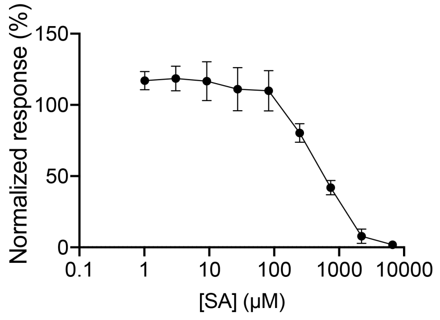


IC$_{50}$ [SA] = 665.5 ± 57.1 µM

**Fig. S7 | Determination of the IC$_{50}$ values for succinylacetone (SA) in parasite growth.** A luciferase-based assay was used for the determination. The IC$_{50}$ values presented in the figure represent means ± SEM of n=5 biological replicates.

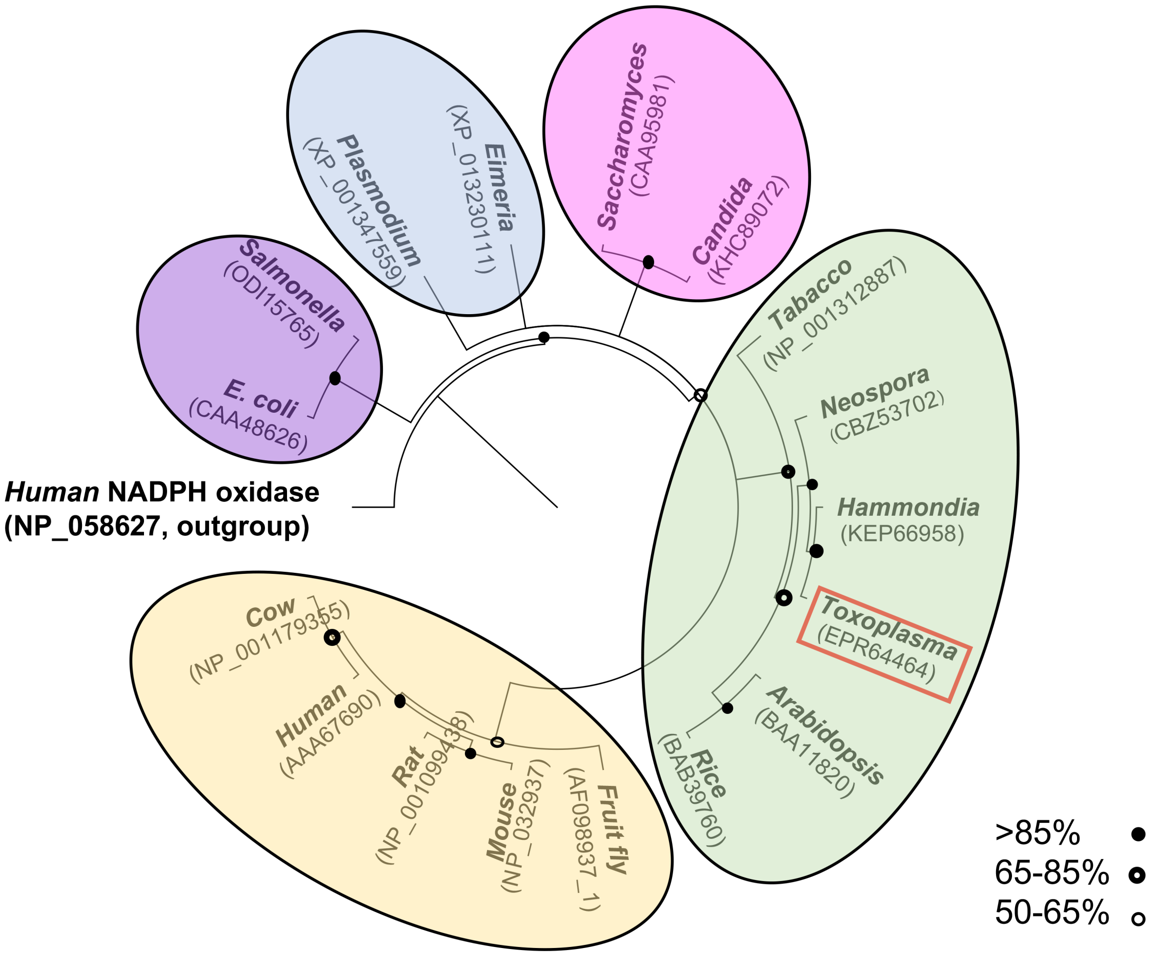


**Fig. S8 | Phylogenetic analysis of protoporphyrinogen oxidase (PPO).** Neighbor-Joining consensus tree analysis of the relationships of 17 PPO family proteins derived from animals, plants, protozoa, fungi, and bacteria. Bootstrap values based on 10000 replicates are shown. Accession numbers of protoporphyrinogen oxidase of individual species were listed in the parentheses. A human NADPH oxidase was also included as an outgroup for phylogeny construction. The closely related PPO orthologs were shaded in individual colors. Consensus bootstrap support (%) was labeled in the figure.

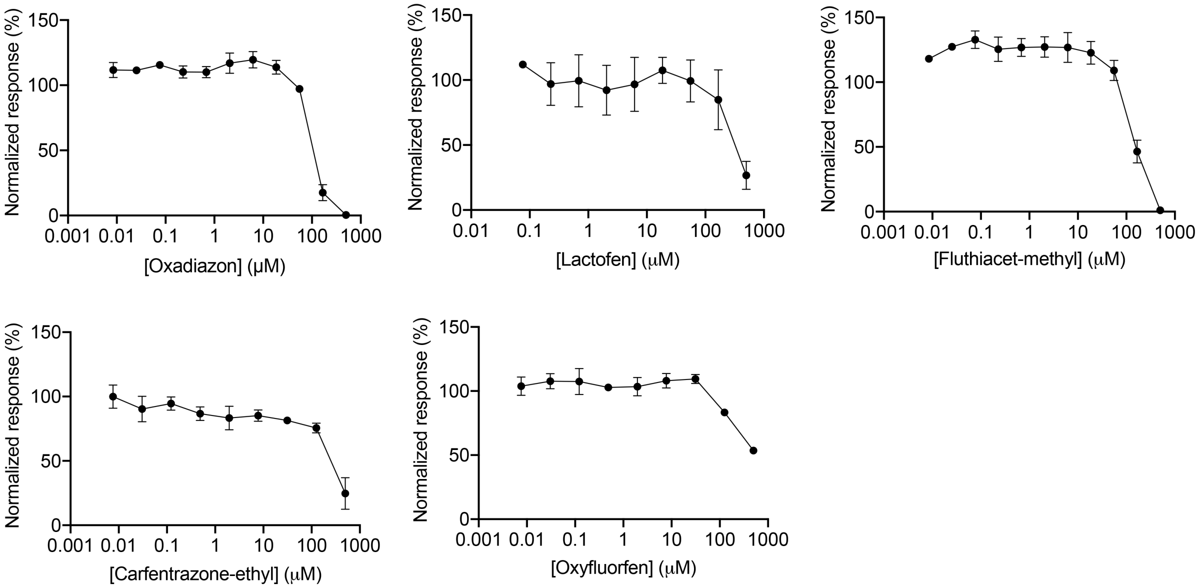


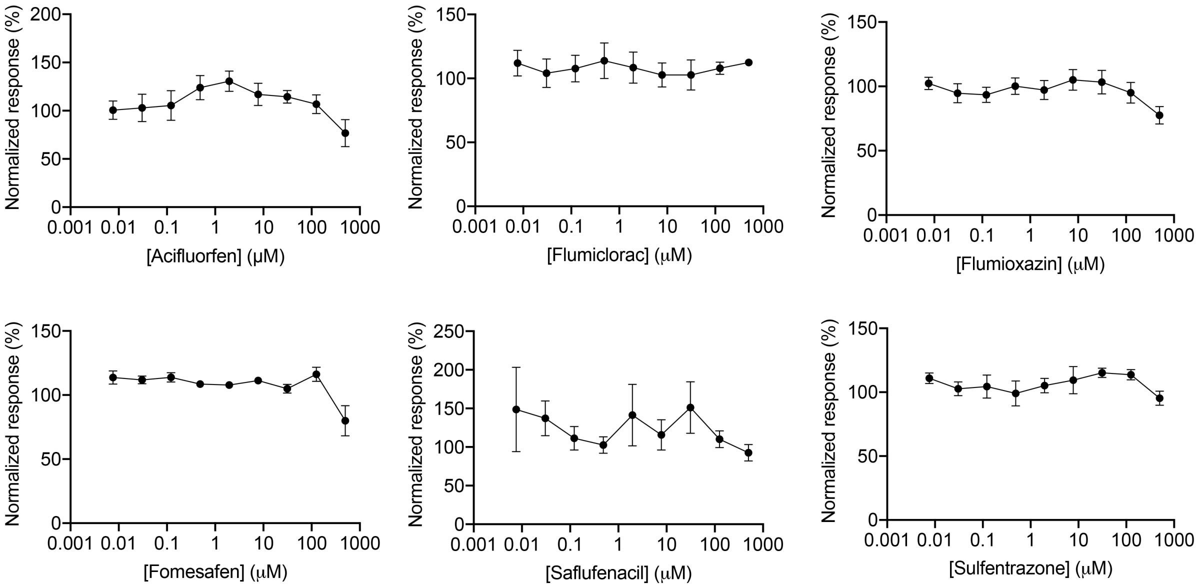


**Table 3. IC50s of the top 5 known PPO inhibitors.**

| | $IC_{50}$ (μM) (mean ± SEM) |
|---|---|
| Oxadiazon | 131.4 ± 3.9 |
| Lactofen | 189.0 ± 38.9 |
| Fluthiacet-methyl | 214.3 ± 25.8 |
| Carfentrazone-ethyl | 262.3 ± 87.0 |
| Oxyfluorfen | 648.3 ± 35.3 |

**Fig. S9 | Screening of 11 commercially available PPO-targeting herbicides in the growth inhibition of *Toxoplasma* parasites. a,** The five most potent inhibitors were identified with their $IC_{50}$ values in the range of ~130 – 650 μM. **b,** Six inhibitors did not show significant inhibitions in parasite growth. The $IC_{50}$ values were reported as means ± SEM of n=3 biological replicates with 3 technical replicates each.

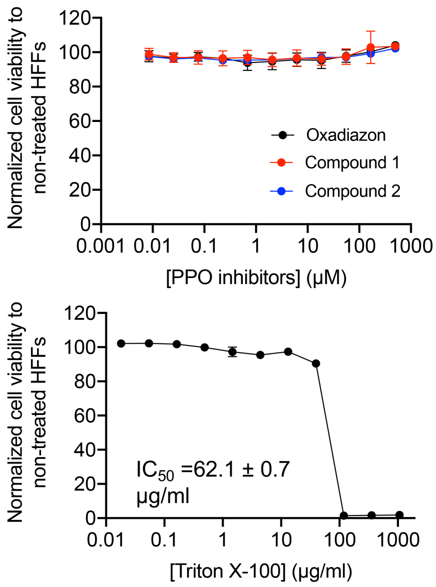


**Fig. S10 | Evaluation of the toxicity of the synthesized oxadiazon derivatives for HFFs.** An AlarmarBlue-based cell viability assay was used to evaluate the toxicity of oxadiazon and its derivatives. Triton X-100 was used as a positive control in the assay. Data represent means ± SEM of n=3 biological replicates with 3 technical replicates each.

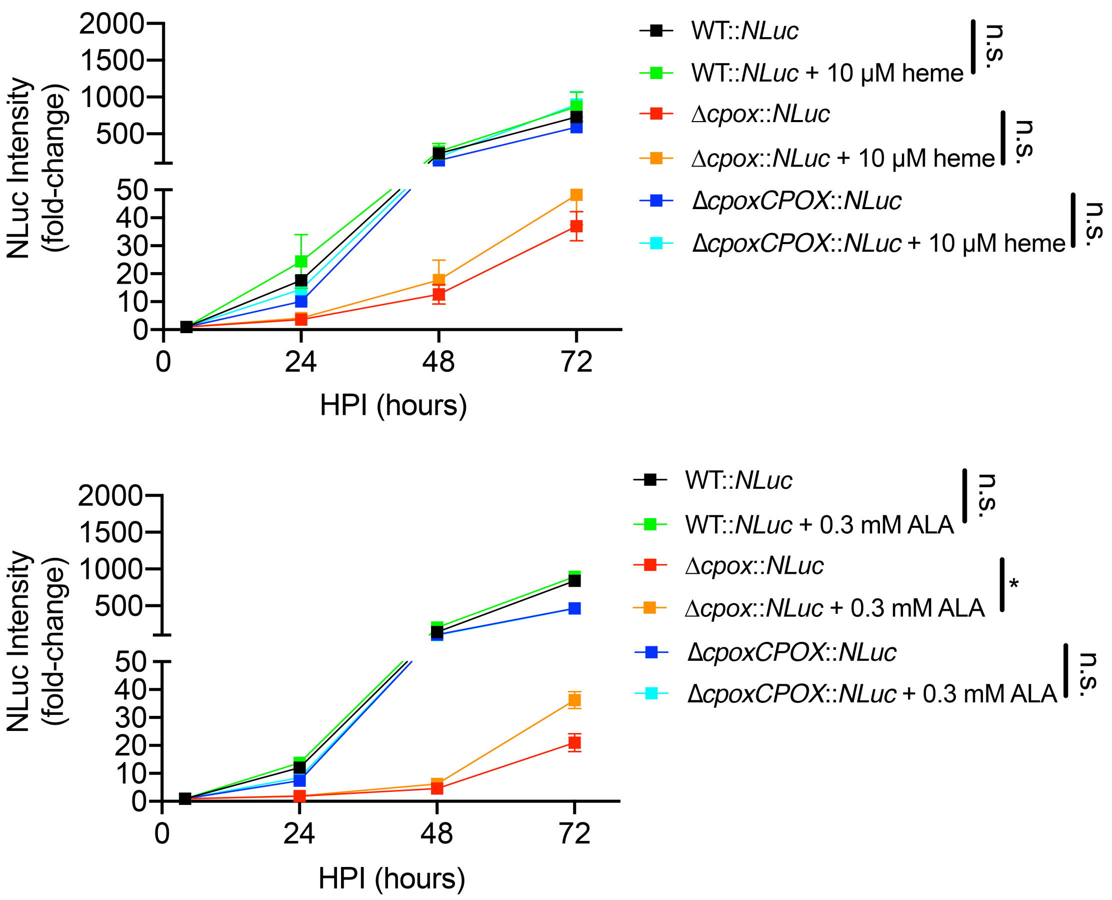


**Fig. S11 | Extracellular heme did not rescue the intracellular growth of the Δ*cpox* parasites.** A luciferase-based growth assay was used to measure the growth of the Δ*cpox* parasites in the media containing or lacking 10 μM heme. The ALA-containing medium was used as a positive control. Data shown here represent means ± SEM of n=3 biological replicates with 3 technical replicates each. Statistical significance was determined by unpaired Student's *t*-test. *, $p<0.05$; n.s., not significant.

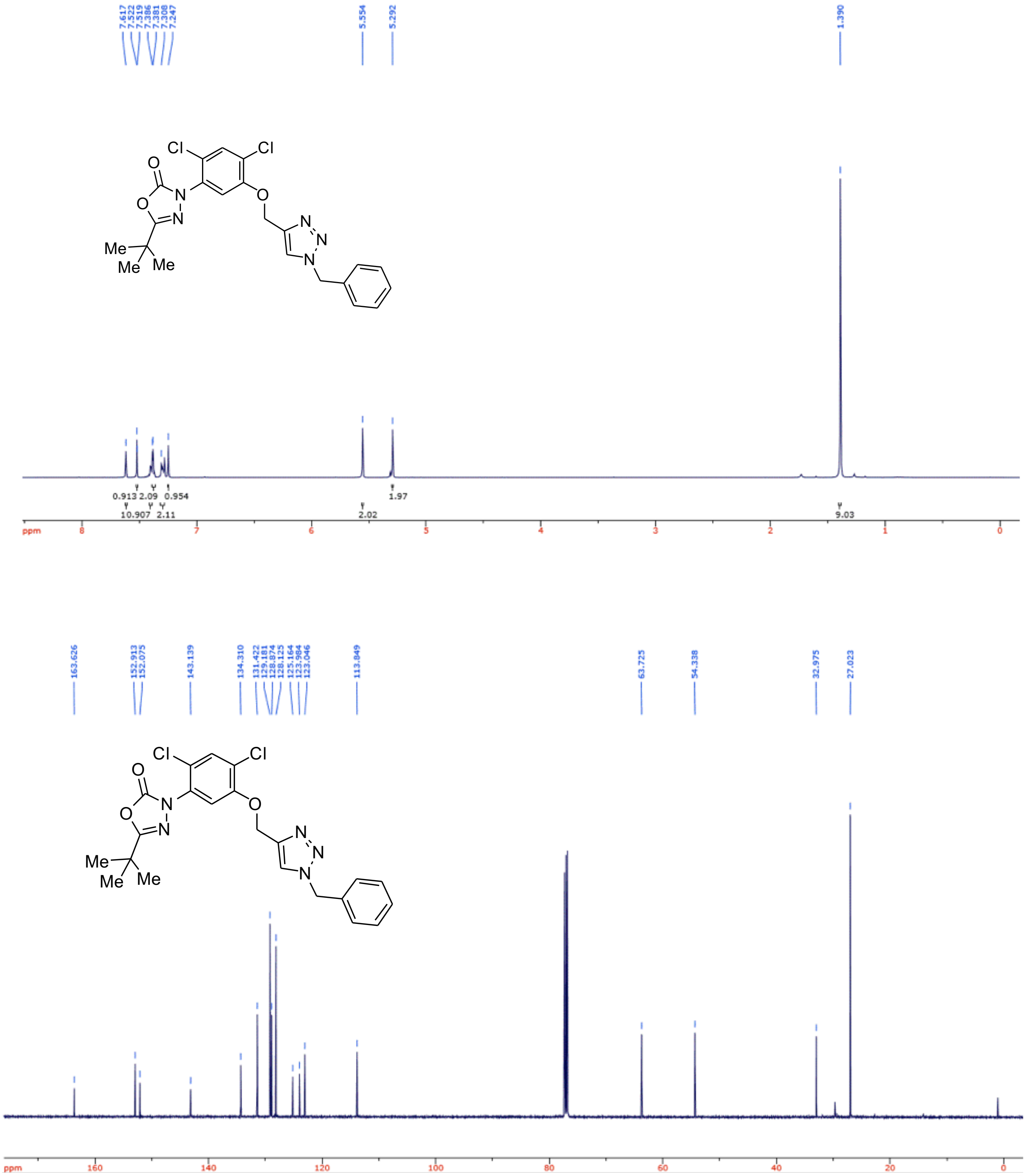


**Fig. S12 | Chemical structure validation of $^{1}$H and $^{13}$C NMR spectra for 3-(5-((1-benzyl-*1H*-1,2,3-triazol-4-yl)methoxy)-2,4-dichlorophenyl)-5-(*tert*-butyl)-1,3,4-oxadiazol-2(*3H*)-one (Compound 1) by $^{1}$H and $^{13}$C NMR spectra.**

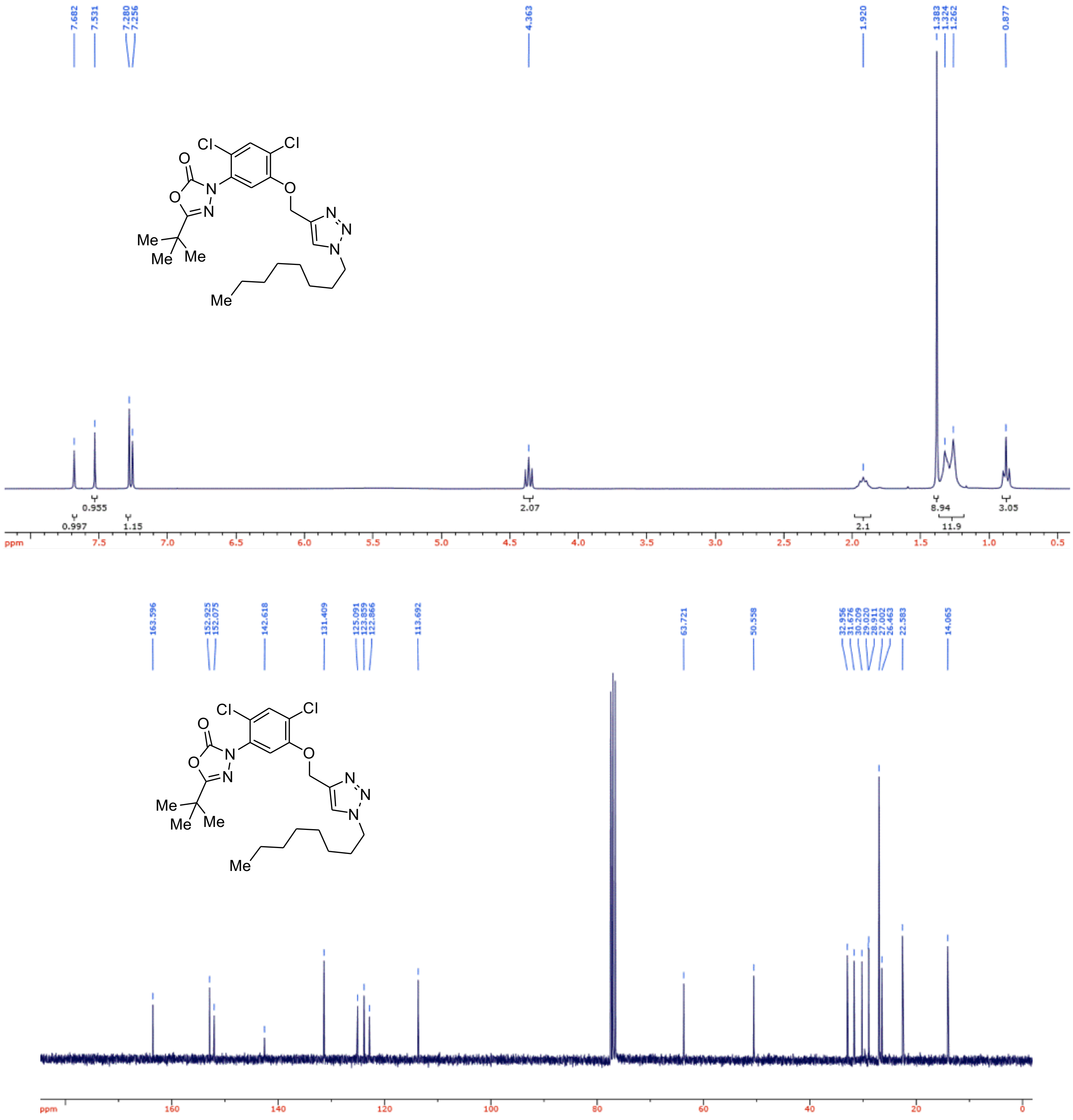


**Fig. S13 | Chemical structure validation of ¹H and ¹³C NMR spectra for 5-(*tert*-butyl)-3-(2,4-dichloro-5-((1-octyl-*1H*-1,2,3-triazol-4-yl)methoxy)phenyl)-1,3,4-oxadiazol-2(*3H*)-one (Compound 2) by ¹H and ¹³C NMR spectra.**
